# Estimating the net effect of functional traits on fitness across species and environments

**DOI:** 10.1101/2022.02.10.479905

**Authors:** Andrew Siefert, Daniel C. Laughlin

## Abstract

1. Functional traits affect the demographic performance of individuals in their environment, leading to fitness differences that scale up to drive population dynamics and community assembly. Understanding the links between traits and fitness is therefore critical for predicting how populations and communities respond to environmental change. However, the net effects of traits on species fitness are largely unknown because we have lacked a framework for estimating fitness across multiple species and environments.
2. We present a modeling framework that integrates trait effects on demographic performance over the life cycles of individuals to estimate the net effect of traits on species fitness. This approach involves 1) modeling trait effects on individual demographic rates (growth, survival, and recruitment) as multidimensional performance surfaces that vary with individual size and environment and 2) integrating these effects into a population model to project population growth rates (i.e., fitness) as a function of traits and environment. We illustrate our approach by estimating performance surfaces and fitness landscapes for trees across a temperature gradient in the eastern United States.
3. Functional traits (wood density, specific leaf area, and maximum height) interacted with individual size and temperature to influence tree growth, survival, and recruitment rates, generating demographic trade-offs and shaping the contours of fitness landscapes. Tall tree species had high survival, growth, and fitness across the temperature gradient. Wood density and specific leaf area had interactive effects on demographic performance, resulting in fitness landscapes with multiple peaks.
4. With this approach it is now possible to empirically estimate the net effect of traits on fitness, leading to improved understanding of the selective forces that drive community assembly and permitting generalizable predictions of population and community dynamics in changing environments.

## INTRODUCTION

Functional traits influence how individuals interact with their abiotic and biotic environment, so that individuals with traits better adapted to their environment will survive and reproduce at greater rates. These trait-based fitness differences scale up to drive population and community dynamics. Quantifying the links between traits and fitness and how they vary across environments is therefore the key to gaining a mechanistic understanding of community assembly and making generalized predictions of population and community dynamics in changing environments (McGill et al., 2006; Shipley et al., 2015). Estimating these trait-fitness relationships has remained largely out of reach, however, due to the empirical challenges of estimating fitness across phenotypes of multiple species, especially for long-lived organisms (Laughlin et al., 2020).

Here, we focus on estimating how trait variation among species affects fitness, defined as the population growth rate of a species in a particular environmental context (McGraw & Caswell, 1996). Population growth is the outcome of the demographic processes of growth, survival, and reproduction over the life cycles of individuals. Establishing links between traits and individual demographic rates is therefore an important first step toward estimating the effect of traits on fitness. Many studies have examined interspecific trait-demographic rate relationships, particularly in trees, but these relationships have often been found to be weak (Yang et al., 2018). A proposed explanation for this is that trait-demographic rate relationships are highly context-dependent (Swenson et al., 2020). The effect of a trait on demographic performance likely depends on an individual’s other traits, size or life stage, and the environment. Recent studies have examined the influence of trait-by-trait, trait-by-size, and trait-by-environment interactions on plant demographic rates across species (Lai et al., 2021; Laughlin et al., 2018; Li et al., 2021), but few if any studies have accounted for all these contexts simultaneously. To do so, we advocate extending the concept of the performance surface— traditionally used by evolutionary biologists to describe the relationship between traits and fitness components across individuals within a population (Arnold, 2003)—to quantify the effects of multidimensional phenotypes on demographic rates across multiple species. Estimating the shape of demographic performance surfaces and how they vary across life stages and environments would bring us a step closer to linking traits with fitness.

Previous studies have quantified the relationship between traits and individual demographic rates, but linking traits to fitness requires integrating trait effects on multiple demographic rates across the life cycle (Laughlin et al., 2020). Single demographic rates are often poor proxies for fitness due to the presence of demographic trade-offs. Traits can generate these trade-offs if they have opposing effects on different aspects of demographic performance, e.g., growth vs. survival (Stearns, 1989). Recent studies have examined demographic trade-offs across many species and examined how species’ positions along these trade-off axes correlate with their functional traits (Adler et al., 2014; Rüger et al., 2018). Directly estimating the effects of traits on multiple demographic rates across multiple life stages would provide stronger direct insights into the role of traits in generating demographic trade-offs that define life-history strategies and determine fitness.

Here we develop a data-driven modeling framework that integrates trait effects on demographic performance over the life cycle to estimate the net effect of traits on fitness across species and environments (Figure 1). To achieve this, we first model the effects of traits on individual demographic rates (growth, survival, and recruitment) as multidimensional performance surfaces whose shape can vary with individual size and local environment, thus accounting for trait-by-size and trait-by-environment interactions. Next, we combine these demographic rate models into a population model (Ellner et al., 2016) that integrates demographic performance across the life cycle to project population growth rates—our measure of fitness—as a function of traits and environment. Using this population model, we estimate the fitness of multidimensional phenotypes in different environments to construct dynamic fitness landscapes that quantify the net effects of traits on fitness across environments. We illustrate our approach by estimating performance surfaces and fitness landscapes for trees across a temperature gradient in the eastern United States and show that trait-mediated demographic trade-offs naturally emerge from the model.

**Figure 1.**
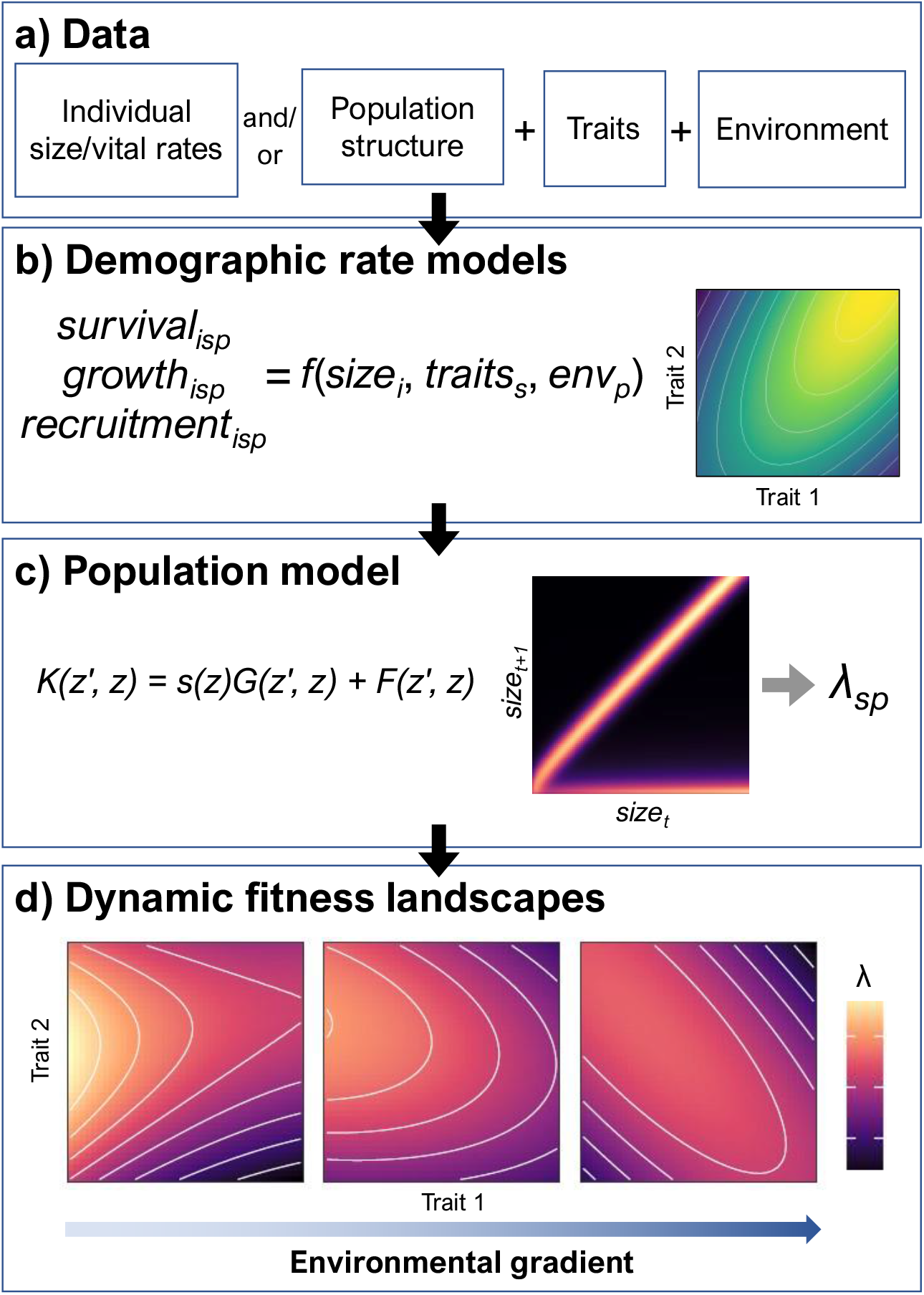
Conceptual illustration of the modeling framework. a) Data on individual sizes and vital rates, species’ traits, and environmental variables are used to b) model individual demographic rates (survival, growth, and recruitment). Population size and structure data (e.g., counts of individuals by size class, number of recruits) can be used in place of or in addition to individual-level data to estimate demographic parameters through inverse modeling approaches. c) The demographic rate models are combined to make a single population model (integral projection model, IPM) that is used to project population growth rates (λ). d) Using the population model, we calculate population growth rates for phenotypes throughout the trait space in different environments to construct dynamic fitness landscapes. Subscripts index individual *i*, species *s*, plot *p*.

## METHODS

### The framework

Previous studies have linked traits with the individual demographic components of fitness— survival, growth, and reproduction—but have not integrated all demographic rates to determine the net effects of traits on fitness (Laughlin et al., 2020). Structured population models integrate demographic performance across the life cycle to project population growth rates, a widely-used measure of population fitness (Caswell, 2001; Ellner et al., 2016). Our approach integrates trait-based demographic models into a single population model that can estimate fitness across species and environments.

The first step in our framework is to model individual growth, survival, and recruitment across species as functions of individual size, species’ traits, and the environment (Figure 1b). Any type of demographic model can be used, but we prefer a Bayesian approach because it allows integration of multiple types of data (Clark et al., 2004), including individual-level and population-level data, as illustrated by our recruitment model in the case study below. Bayesian models also make it easy to quantify uncertainty in demographic parameters and propagate this uncertainty to the population model. Our framework is flexible with respect to the functional forms used to quantify the effects of traits, size, and environment. Drawing inspiration from evolutionary theory (Lande & Arnold, 1983), we model trait effects as multidimensional surfaces that can capture linear and nonlinear effects and trait-by-trait interactions. By allowing the shapes of these surfaces to vary depending on individual size and the environment, this approach also can account for trait-by-size and trait-by-environment interactions.

The second step in our framework is to integrate the demographic rate models into a single trait-based population model (Figure 1c) and use it to estimate the fitness of different phenotypes in different environments. We use integral projection models (IPMs) due to their flexibility. IPMs combine information about individuals’ size-specific survival, growth, and recruitment rates to project population dynamics (Merow, Dahlgren, et al., 2014). An IPM describes how the size distribution of individuals in a population changes through time:

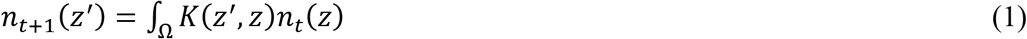

where *n*_*t*_(*z*) is the size distribution at time *t, n*_*t+1*_(*z’*) is the size distribution at time *t* + 1, Ω denotes the possible range of individual sizes, and the kernel *K*(*z’,z*) describes size transitions through survival, growth, and reproduction:

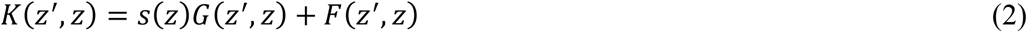

where *s*(*z*) is the survival probability dependent on initial size *z, G*(*z’,z*) describes the probability of growing from size *z* to *z’*, conditional on having survived, and *F*(*z’,z*) describes the size distribution of new recruits produced by an individual of size *z*. These demographic transitions can be estimated using the trait-based demographic rate models described in step 1, allowing construction of IPM kernels for hypothetical populations with any combination of trait values (i.e., phenotype) in any environment. The long-term population growth rate (λ) can be extracted by eigenanalysis of the discretized kernel. λ is the growth rate to which a population will converge if its demographic rates remain constant and it is allowed to reach its stable size distribution. It provides an integrative relative measure of population performance, summarizing how all demographic processes over the life cycle combine to determine how fast a population grows (Ellner et al., 2016), and is commonly used in evolutionary biology as a measure of fitness. Transient growth rates—the expected short-term growth rate of a population given its current size distribution and demographic rates—can also be calculated by using the kernel to project the population forward in time (Merow, Dahlgren, et al., 2014). We quantify fitness landscapes by constructing IPM kernels and extracting long-term population growth rates for phenotypes throughout the trait space in different environments (Figure 1d).

It is important to note that these fitness landscapes describe the mapping from phenotype to fitness irrespective of species identity. If there are interspecific differences in demographic rates not explained by traits (e.g., as captured by species random effects in the demographic rate models), this information could be included when constructing the IPM kernels to provide species-specific fitness estimates.

### Case study: fitness landscapes for temperate trees

We illustrate our approach by estimating fitness landscapes for trees across a temperature gradient in the eastern United States. We fit demographic models using data from the US Forest Service Forest Inventory and Analysis (FIA; http://www.fia.fs.fed.us). The FIA dataset consists of forest plots that are censused at varying intervals. At each census, individual survival and diameter growth are measured for all trees ≥12.7 cm diameter at breast height (dbh; “canopy trees” hereafter) within a plot and all saplings (2.54-12.7 cm dbh) within smaller microplots. We extracted data from 12,752 plots in the eastern United States censused between 2003 and 2019 (see the Supplement for details about FIA plot design and data selection). We extracted mean annual temperature data for the years spanning the census interval for each plot from gridMET (Abatzoglou, 2013).

We focused on three traits representing key axes of plant functional strategies: wood density, specific leaf area (SLA), and maximum height. Wood density reflects trade-offs between stem hydraulic efficiency, hydraulic safety, and mechanical strength (Chave et al., 2009). Specific leaf area reflects the trade-off between the cost of leaf construction and rate of return on investment in carbon and nutrients (Wright et al., 2004). Maximum height reflects a trade-off where taller species are better competitors for light but have higher stem construction and maintenance costs and deferred reproduction (Falster & Westoby, 2003). We extracted trait values for tree species in FIA plots from the TRY plant trait database (Kattge et al., 2020).

Survival and growth were modeled at the individual level. We created separate survival models for saplings and canopy trees because size effects on survival across all sizes were not well described by several functional forms we tried. We modeled growth of saplings and canopy trees together, using average annual diameter growth rate as the response variable. Recruitment was measured at the plot level as the number of trees crossing the 2.54-cm threshold during the census interval (i.e., ingrowth). For each data set (sapling survival, canopy tree survival, growth, and recruitment), we split the data into a training set (80% of plots) for model fitting and test set (20% of plots) for model evaluation. We excluded plots containing fewer than 10 individuals (5 individuals for sapling survival) and species occurring in fewer than 10 plots (5 plots for sapling survival). The final training data sets contained: 224,153 trees, 8,837 plots, 94 species (canopy tree survival); 45,249 trees, 4,518 plots, 78 species (sapling survival); 250,768 trees, 9,152 plots, 95 species (growth); 32,891 plot-species observations, 4,099 plots, 83 species (recruitment).

To improve estimates of the relationship between tree size and recruitment, we obtained data on individual size and reproductive status (presence of reproductive structures) from the MASTIF network (Clark et al., 2019). We selected data from sites in eastern North America that had at least one species in common with our demography modeling data set. Data were collected between 2002 and 2020, with some trees being measured in multiple years. We excluded observations for which reproductive status was unknown, resulting in a data set of 48,082 observations of 27,641 trees from 60 species in 34 sites.

One concern about estimating demographic performance using observational data is that if we only observe species in favorable environments where they can persist (λ ≥ 1), we will lack information about how performance varies across environments. We think it is likely, however, that large observational data sets such as FIA include observations of species in both favorable and unfavorable environments and so include demographic failures (λ < 1). To test this, we fit population models for four representative species with ample FIA and MASTIF data and estimated their population growth rates across the ranges of mean annual temperatures in which they occurred. We found that λ varied considerably with mean annual temperature for each species, including values above and below 1 (see Supporting Information Figure S10), confirming that our data set captured variation in demographic performance across the temperature gradient.

#### Demographic rate models

We modeled survival, growth, and recruitment using hierarchical Bayesian models that included terms representing the effects of size, crowding, climate, and traits, as well as species and plot random effects. Here we present an overview of salient features of the models. Additional modeling details are included in the Supporting Information, and descriptions of all terms and parameters in the demographic rate models are provided in Tables S1-S4.

Trait effects on demographic rates were modeled as multivariate Gaussian surfaces (i.e., performance surfaces; Lande 1980) whose shape varied with temperature:

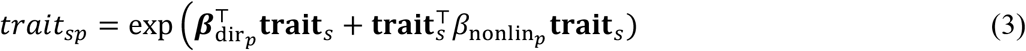

where *trait*_*sp*_ is the effect of traits on the demographic performance (survival, growth, or reproduction) of species *s* in plot *p*, 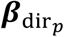 is a vector of directional (linear) performance gradients in plot *p*, and 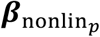 is a matrix of nonlinear performance gradients in plot *p* (Arnold, 2003). The diagonal elements of ***β***_nonlin_ measure the strength of stabilizing (if *β* is negative) or disruptive (if *β* is positive) selection for each trait, and the off-diagonal elements measure the strength of correlational selection between trait pairs (Arnold, 2003). Positive correlational selection means that performance is maximized by having either high or low values of both traits. Negative correlational selection means that performance is maximized by having a high value of one trait and low value of the other trait. This function can produce performance surfaces of various shapes, including (when viewed in 2 dimensions) a peak, a saddle, a ridge, or a slope.

To allow performances surfaces to vary across the temperature gradient, the performance surface parameters (elements of ***β***_dir_ and ***β***_nonlin_) were modeled as linear functions of mean annual temperature:

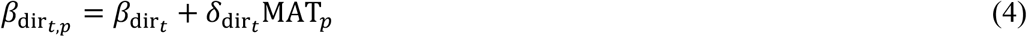

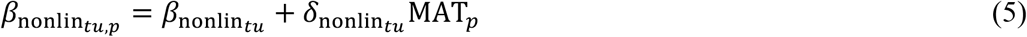

where 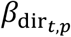 is the directional selection coefficient for trait *t* in plot *p*, 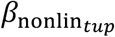 is the stabilizing/disruptive selection coefficient for trait *t* (if *t* = *u*) or correlational selection coefficient for traits *t* and *u* (if *t* ≠ *u*) in plot *p*. 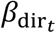 and 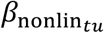 are the performance gradients in a plot with average temperature, and 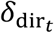 and 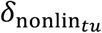 describe how the trait effects change along the temperature gradient (i.e., trait-by-environment interactions).

The effect of tree size on demographic performance was modeled as an increasing (sapling survival and recruitment) or hump-shaped (canopy tree survival and growth) function of individual diameter. The parameters of the size functions were themselves functions of traits, allowing for trait-by-size interactions. The effect of crowding was modeled as a decreasing (power law) function of the total basal area of neighboring trees (see the Supplementary Methods for additional details).

Whereas individual survival and growth could be modeled directly using individual-level data, reproduction was modeled using an inverse approach that combined information on plot-level recruitment and individual-level reproductive status (Clark et al., 2019). Briefly, for each tree in a plot, the model estimated the annual production of recruits as a function of size, crowding, climate, and traits. The size effect was a product of the probability of being reproductive and the per-capita production of recruits, conditional on being reproductive (Ribbens et al., 1994). These annual per-capita recruitment rates were summed across trees in a plot and multiplied by the census interval (in years) to give the predicted number of recruits. By modeling plot-level recruitment (using FIA data) and individual-level reproductive status (using MASTIF data) jointly within a Bayesian framework, both types of data informed estimates of individual recruitment rates.

We assessed model performance by calculating their predictive accuracy (AUC for survival, *R*^2^ for growth and recruitment) on the training and test sets. To assess the ability of traits to predict species’ demographic rates, we calculated the proportion of variation in species’ expected demographic rates (i.e., predictions including species random effects) explained by traits (i.e., predictions based on species’ traits only, excluding species random effects) within different size and temperature bins.

#### Trait-mediated demographic tradeoffs

Life history theory holds that organisms face trade-offs in allocation to different demographic processes across the life cycle, resulting in demographic trade-offs that constrain life history strategies (Stearns, 1989). Key demographic trade-offs posited for trees include the growth-survival trade-off, which is thought to be strongest at the sapling stage (Wright et al., 2010), and the stature-recruitment trade-off, which distinguishes between phenotypes that recruit early in life versus those that exhibit high growth and survival later in life (Rüger et al., 2018). We explored whether these trade-offs emerged from the effects of functional traits on demographic rates in our model. To test the role of traits in mediating a sapling growth-survival trade-off, we calculated and plotted the predicted growth vs. survival of trees at 5 cm dbh across a range of trait values, with other model predictors held constant at their average values. To test whether traits generated a stature-recruitment trade-off, we similarly plotted the predicted growth or survival at 60 cm dbh vs. recruitment at 8 cm dbh (the average size at onset of reproduction across all species) across a range of trait values.

#### Fitness landscapes

To construct fitness landscapes, we integrated the demographic rate models described above into a population model (IPM) that we used to project population growth rates of multidimensional phenotypes across the temperature gradient. We constructed IPM kernels and extracted long-term population growth rates (λ) for a grid of trait values covering the observed trait space at each of three mean annual temperatures (5, 10, and 15°C). Further details about IPM implementation are provided in the Supporting Information.

## RESULTS

### Trait-demographic rate relationships

Functional traits influenced tree survival, growth, and recruitment rates, and these effects varied with tree size and mean annual temperature (see Tables S6-S9 for parameter estimates and credible intervals). The sapling and canopy tree survival models had AUC of 0.79 and 0.77, respectively, on the training set and 0.69 on the test set (Fig. S11b,c,e,f). Traits explained 8-75% of variation in species’ survival rates, depending on the size class and mean annual temperature, with more variation explained for larger-diameter trees (Fig. S12a). Tree species with the tallest maximum heights and densest wood had the highest survival rates (Figure 2a-d, Figure S2a,c). Wood density and SLA had an interactive effect on survival, especially of small-diameter trees, such that survival peaked for species with either high wood density and high SLA or low wood density and low SLA (Figure 2a, Figure S4a).

**Figure 2.**
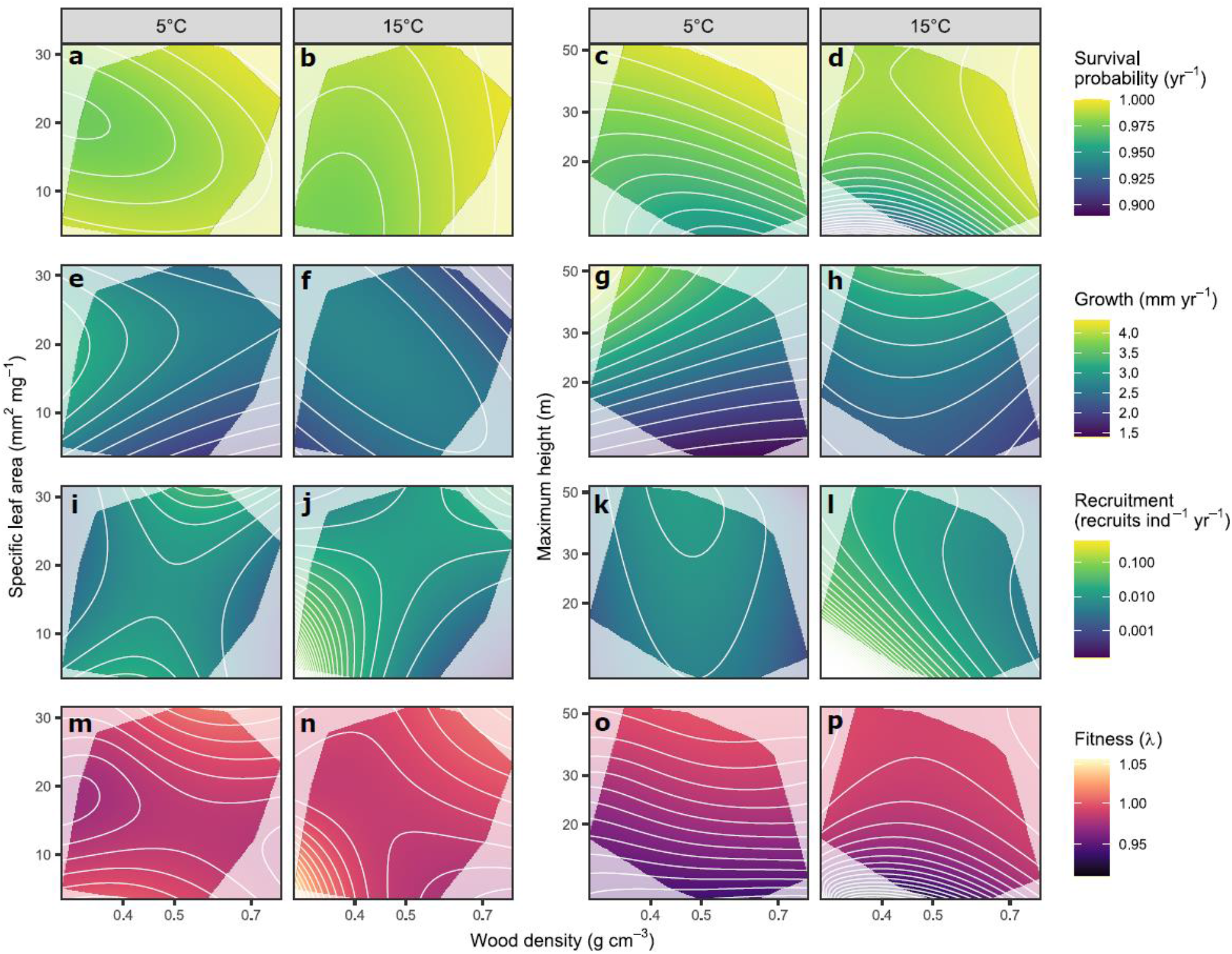
Tree performance and fitness landscapes at low and high mean annual temperatures. Landscapes show expected demographic rates (survival, a-d; diameter growth, e-h; recruitment, i-l) and fitness (population growth rate, m-p) for trees species with different trait combinations. Demographic rate and fitness models included three traits—wood density, specific leaf area (SLA), and maximum height—but for ease of visualization, landscapes are shown for two traits at a time (wood density and SLA, columns 1-2; wood density and maximum height, columns 3-4) with the third trait held constant at its average value. Demographic rates also vary with tree size, mean annual temperature, and neighbor density in our model. Performance landscapes shown here are for trees with 20 cm diameter at low (5°C, columns 1 and 3) or high (15°C, columns 2 and 4) mean annual temperature and average neighbor density. Fitness landscapes integrate demographic performance across sizes. Grayed areas show regions of trait space not occupied by tree species in our data set.

The growth model explained 35% and 23% of variation in individual tree growth rates in the training and test sets, respectively (Fig. S11h,i). Traits explained 12-36% of variation in species’ growth rates depending on size and mean annual temperature, with more variation explained for large-diameter trees (Fig. S12b). Species with the tallest maximum heights and lowest wood densities grew the fastest, especially in cold sites (Figure 2g,h, Figure S2d,f). There was a complex interaction between wood density, SLA, and temperature. In cold sites, growth peaked at low values of wood density and medium to high values of SLA (Figure 2e), whereas in warm sites there was a ridge of high growth rates in the performance surface running from an acquisitive strategy of low wood density and high SLA to a conservative strategy of high wood density and low SLA (Figure 2f, Figure S5a). The positive effect of SLA on growth was strongest in small-diameter trees (Figure S2e). In contrast, the effect of maximum height was strongest for large-diameter trees (Figure S2f).

The full recruitment model explained 36% and 13% of variation in population-level recruitment in the training and test sets, respectively (Fig. S11k-l). Traits explained 5-67% of variation in species’ recruitment rates depending on size and temperature, with more variation explained for larger-diameter trees and warmer sites (Fig. S12c). Species with low wood density had the highest per-capita recruitment rates, particularly in warm sites (Figure 2j,l, Figure S3a). There was strong positive correlational selection between wood density and SLA, such that recruitment peaked at a combination of low wood density and low SLA and a combination of high wood density and high SLA, especially in warm sites (Figure 2j, Figure S6a). The effects of maximum height and wood density on per-capita recruitment depended on tree diameter. For small-diameter trees, per-capita recruitment was strongly negatively related to wood density and maximum height, whereas for large-diameter trees these effects were weaker (Figure S3a,c). These interactions occurred because species with dense wood and tall maximum height had a larger size at onset of reproduction (Figure 3d).

**Figure 3.**
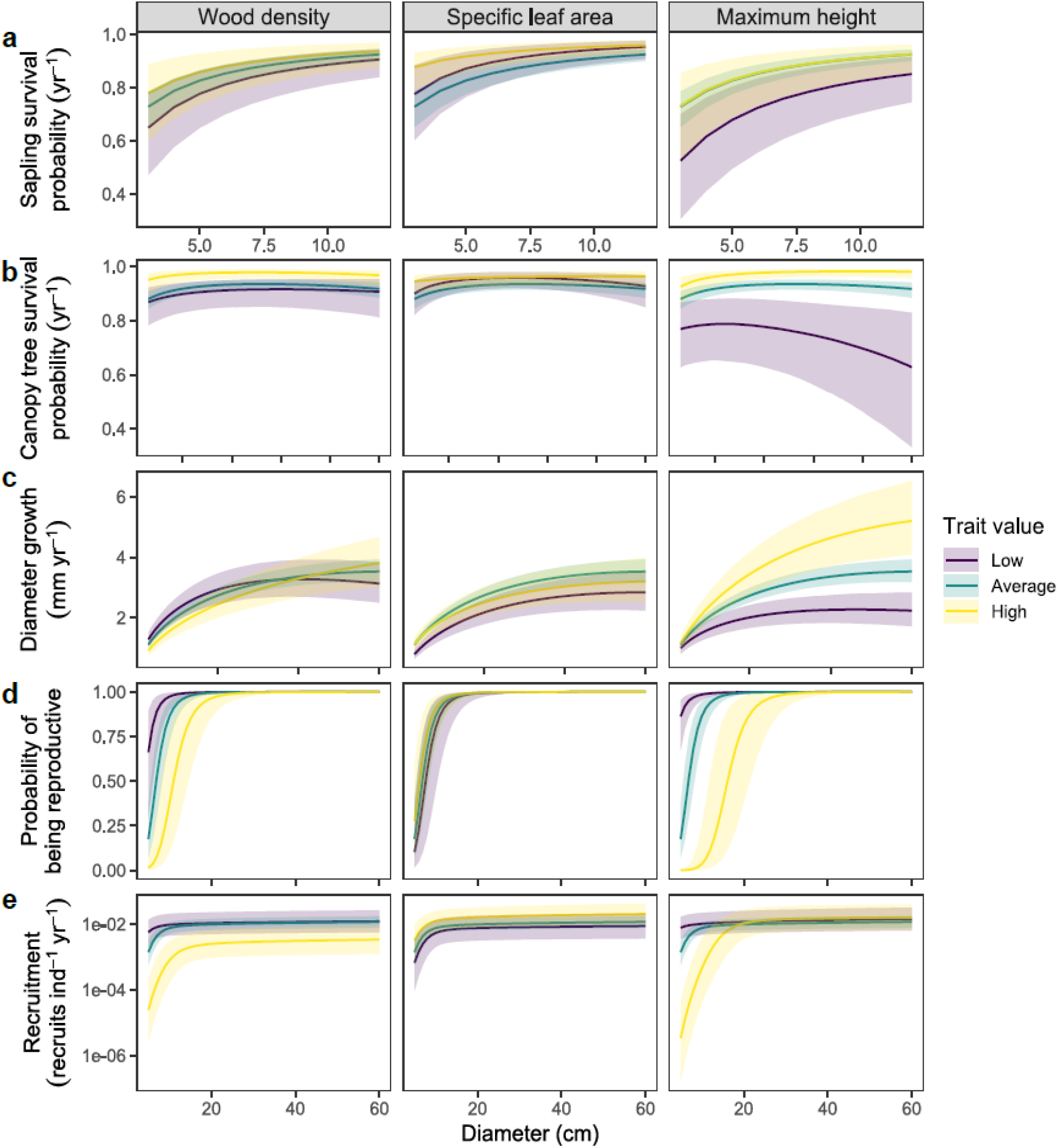
Relationship between tree diameter and demographic performance with respect to functional traits. Plots show the relationship between tree diameter and expected sapling survival (a), canopy tree survival (b), diameter growth (c), probability of being reproductive (d), and annual per-capita recruitment (e). Trend lines show conditional expectations for trees with different values of wood density (column 1), specific leaf area (column 2), or maximum height (column 3) with other traits, temperature, and neighbor density held constant at their average values. Size-demographic rate relationships are show for average, low (average – 2SD), and high (average + 2SD) values of each trait to illustrate trait-by-size interactions. Shaded areas show 90% credible intervals.

### Trait-mediated demographic trade-offs

We found evidence for a growth-survival trade-off among saplings driven by wood density. Saplings of species with low wood density grew quickly but had low survival rates, whereas saplings of species with high wood density had high survival rates but grew slowly (Figure 4a). This growth-survival trade-off naturally emerged from the opposing effects of wood density on growth and survival in our models. We also found evidence of a stature-recruitment trade-off mediated by maximum height and wood density. Species with low maximum height and low wood density had high recruitment at small sizes but low growth and survival at larger sizes, whereas species with tall maximum height and dense wood had high growth and survival as large trees but produced few recruits when they were small (Figure 4b,c,h,i). We did not find evidence of a growth-survival trade-off or stature-recruitment trade-off driven by specific leaf area (Figure 4d-f).

**Figure 4.**
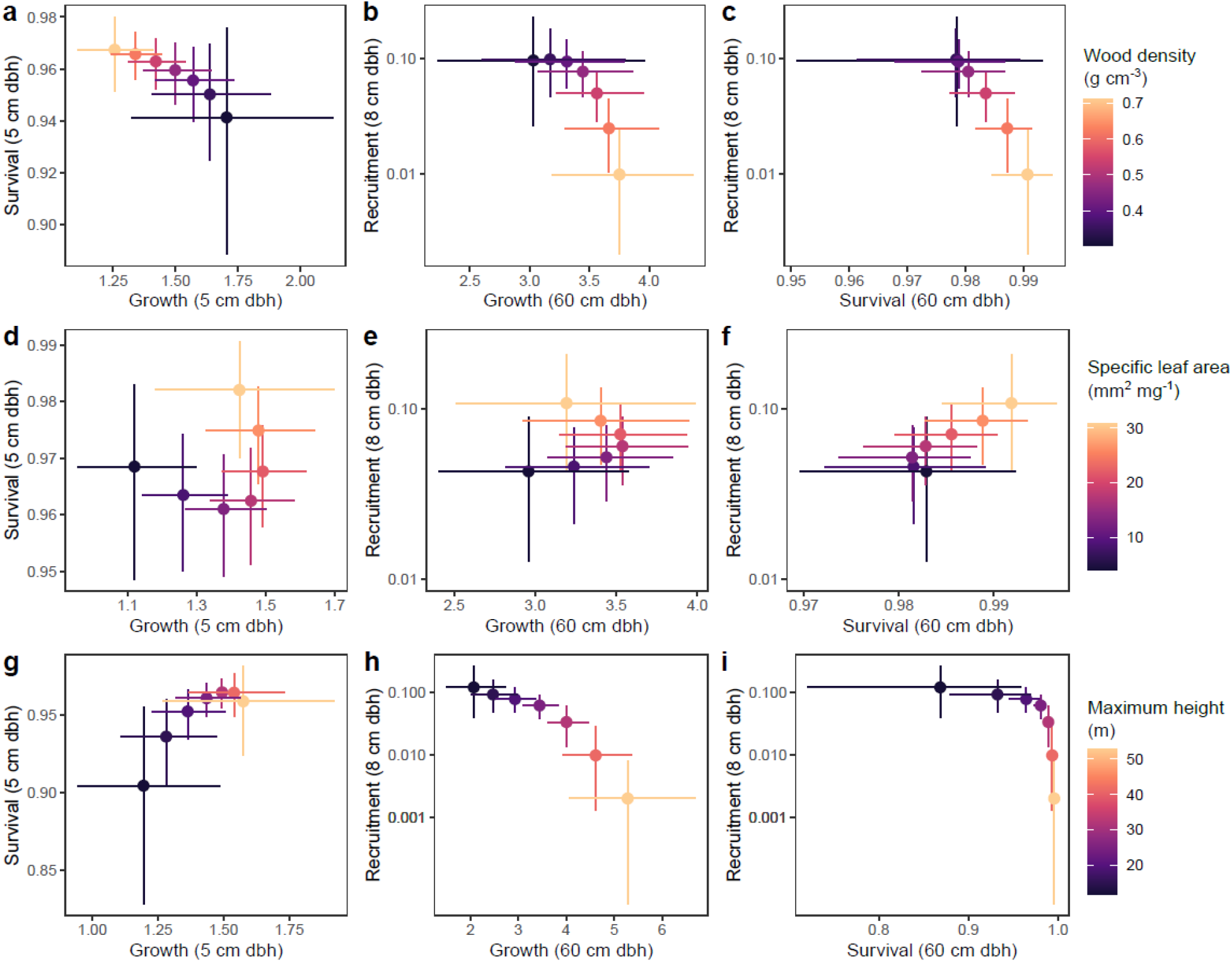
Trait-mediated demographic trade-offs. Points represent predicted demographic rates (error bars show 90% credible intervals) for trees with varying wood density (a-c), specific leaf area (d-f), or maximum height (g-i), with other model predictors held at their average values. A negative relationship between demographic rates across values of a trait indicates a trait-mediated demographic trade-off. The first column (a,d,b) shows the relationship between predicted growth and survival at 5 cm diameter, diagnostic of a growth-survival tradeoff in saplings. The second (b,e,h) and third (c,f,i) columns show relationships between growth and survival, respectively, at 60 cm diameter and recruitment at 8 cm diameter, diagnostic of a trade-off between recruitment early in life and growth and survival later in life (i.e., stature-recruitment tradeoff).

### Fitness landscapes

Tree species with taller maximum heights had higher fitness at all temperatures, especially in colder sites (Figure 2o-p, Figures S7b,c, S8). The positive overall effect of maximum height on fitness likely resulted from its positive effects on both growth and survival across the life cycle (Figure S2c,f). The net effects of wood density and specific leaf area on fitness were weaker, nonlinear, interactive, and variable across the temperature gradient. The strongest effects of wood density and SLA were in warm sites, where fitness was highest for species with low wood density and low SLA (Figure 2n, Figure S7a), likely due to these species having high sapling survival coupled with high recruitment rates (Figures S4a, S6a). Fitness landscapes for wood density and SLA were bimodal, with fitness peaks at both low wood density coupled with low SLA and high wood density coupled with high SLA (Figure 2m-n, Figure S7a). This appears to be driven by a similar bimodal shape of the performance landscape for sapling survival (Figure S4a). These bimodal and relatively flat fitness landscapes indicate the presence of alternative functional strategies that yield similar fitness.

## DISCUSSION

The idea that traits drive variation in species fitness is a core assumption of trait-based ecology (Violle et al., 2007) and foundational to the promise that trait-based approaches can make community ecology more general, mechanistic, and predictive (Lavorel & Garnier, 2002; McGill et al., 2006; Shipley et al., 2015). Despite progress in estimating the relationship between traits and the demographic components of fitness, we still lack information about the net effects of traits on fitness itself, due in large part to the empirical challenges of estimating fitness across phenotypes of multiple species in multiple environments. Our approach overcomes these challenges by integrating trait-based demographic models into a single population model to project population fitness as a function of multidimensional phenotypes and the environment. With this approach it is now possible to empirically estimate the net effect of traits on fitness, leading to improved understanding of the selective forces that drive community assembly and permitting mechanistic predictions of community dynamics in changing environments.

### Insights into the adaptive value of traits

Community ecologists have long been interested in understanding how traits drive the performance and distributions of species across environmental gradients. The metaphor of an environmental filter is often used to describe natural selection acting across populations within a community, whereby species with certain trait values succeed and persist in an environment and others fail (Keddy, 1992). Previous work has sought to infer environmental filters by examining trait-environment relationships and trait distribution patterns (e.g., Cornwell & Ackerly, 2009; Laughlin et al., 2012). However, these patterns reflect the aggregated effects of multiple processes, including selection, dispersal, and ecological drift, acting across multiple generations (Lasky et al., 2013). Our approach translates the metaphor of environmental filtering into an operational framework to directly estimate the effects of traits on performance and the likelihood of population persistence. Quantifying these effects is critical for understanding the adaptive value of traits in different environments and predicting how populations and communities respond to environmental change.

Our case study shows how our modeling framework can provide insights into the adaptive value of functional traits within communities. We found that functional traits influenced the demographic performance and fitness of trees across the eastern United States. Tall species had relatively high survival and grew quickly throughout their life cycle but produced few recruits at small sizes, reflecting a strategy of “long-lived pioneers” previously described in tropical forests (Rüger et al., 2018). Short species reached reproductive maturity quickly and had high recruitment rates at small diameters, but this had relatively little overall fitness benefit because fitness of long-lived plants is more sensitive to survival and growth than to reproduction (Franco & Silvertown, 2004). The low fitness of short-statured species (λ < 1, indicating population decline) might seem surprising, but it likely reflects the fact that we evaluated fitness at the average crowding level (i.e., total stand basal area) in our data set, which only included forests at least 20 years old. Many short-statured species are shade intolerant, early-successional species that are likely outcompeted by taller, late-successional species by this time (Falster & Westoby, 2005). Some short tree species are able to persist in the understories of mature forests, so their low predicted fitness suggests that our model did not adequately account for processes, such as gap formation and spatio-temporal niche partitioning, that enable their persistence (Falster et al., 2017). However, these temporal dynamics could be integrated and estimated provided that enough empirical observations in young, regenerating forests stands are available.

Wood density and specific leaf area had interactive effects on demographic rates, and these effects varied over the life cycle and across the temperature gradient, exemplifying the context-dependence of trait-demographic rate relationships (Yang et al., 2018). For example, for large-diameter trees in cold sites, species with low wood density and high SLA grew the fastest, consistent with the expectation that these trait values are part of a fast, resource-acquisitive strategy (Reich, 2014). In contrast, for large-diameter trees in warm sites, species with the opposite trait values (high wood density and low SLA) also grew fast, indicating that a conservative strategy might be beneficial for growth given the hydraulic challenges faced by large trees at warm temperatures (Fajardo et al., 2019). Indeed, species with these trait values, including southern oaks (e.g., *Quercus nigra, Q. falcata*) and hard-wooded pines (e.g., *Pinus taeda, P. elliotti*) are known to be fast-growing in the southern US (Burns & Honkala, 1990). Wood density and specific leaf area also had interactive effects on sapling survival and recruitment (which integrates seed production and seedling performance, because we counted recruits as trees reaching the 1-cm diameter threshold), producing bimodal performance surfaces in which performance peaked for species with either high wood density and high SLA or low wood density and low SLA. These effects produced similarly bimodal fitness landscapes, providing evidence of alternative functional strategies that have similar fitness, potentially contributing to the maintenance of functional diversity (Marks & Lechowicz, 2006).

### Predicting population and community dynamics

Predicting the responses of a large number of species to climate change and other global change drivers is an ongoing challenge for ecologists. Demographic population models have been used to project population dynamics for single species across different environments (Merow, Latimer, et al., 2014), but measuring demographic rates for every species in every environment is not feasible. Our approach of modeling population growth rates as a function of traits and environment allows for generalization to species with known traits but limited demographic data and to populations in environments outside their species’ current realized niche (Butt & Gallagher, 2018; Evans et al., 2016).

Although our case study focused on estimating long-term population growth rates, the trait-based population models (IPMs) we describe can also estimate transient population growth rates and incorporate environmental and demographic stochasticity (Ellner et al., 2016). These models can therefore be used to simulate population dynamics in a changing environment by 1) calculating demographic rates as a function of the environment at a specific time point, 2) constructing an IPM kernel using those environment-specific demographic rates, 3) using the kernel to project the population forward to the next time point, and 4) repeating through time.

This approach can be extended to simulate community dynamics by simultaneously projecting the dynamics of multiple interacting species. This is essentially the approach used by forest simulation models (Botkin et al., 1972; Strigul et al., 2008), which simulate forest dynamics based on the demography of interacting individuals or cohorts. A key limitation of these models is that they are difficult to parameterize for many species. As a result, species are often grouped into broad functional types, and some demographic parameters, particularly recruitment, are typically assumed to be fixed across all species (Moorcroft et al., 2001; Purves et al., 2008). Our approach using species’ traits to inform estimates of demographic parameters can help overcome this limitation and make these models more generalizable.

### Other extensions and limitations

Given the flexibility of our framework, it can be adapted and extended in many ways. Here we present a few examples. First, although we modeled the effect of the biotic neighborhood as a simple function of total neighbor abundance, more complex forms of density and frequency dependence could be included. For example, size-structured competition for light is a common feature of forest models (Pacala et al., 1996; Strigul et al., 2008) and could be incorporated into the demographic rate models in our framework. Responses to competition could also be modeled as a function of the traits of both the target population and its neighbors (Kunstler et al., 2012), allowing exploration of how frequency-dependent interactions warp the fitness landscape.

Crucially, stable coexistence requires that species limit themselves more strongly than they limit their competitors, allowing species to increase when rare, i.e., invade a resident community (Chesson, 2000). Our framework could be used to calculate invasion growth rates and partition the contribution of trait differences according to modern coexistence theory (Ellner et al., 2019), providing insights into the role of functional traits in maintaining species diversity.

Second, whereas we used a fixed mean trait value for each species, the models could incorporate intraspecific trait variation. Functional traits can vary strongly within and among populations within species, and this variation can affect demographic performance (Bolnick et al., 2011). The simplest way to incorporate intraspecific trait variation in our framework would be to replace overall species mean trait values with site-specific species mean trait values. This introduces greater data requirements, but modeling traits themselves as a function of the environment would allow estimation of site-specific trait values without the need to measure traits in every site. Intraspecific trait variation within sites could also be incorporated by treating different phenotypes as distinct “populations” and modeling their dynamics separately, or by including traits as additional state variables (i.e., in addition to size) in the IPMs (Ellner et al., 2016).

Finally, all the analytical tools developed for matrix population models, including life table and perturbation analysis, can be applied to the trait-based population models in our framework (Caswell, 2001). For example, the models can be used to calculate life history traits, such as life expectancy and age at reproductive maturity, and examine how they vary with functional traits and the environment. Exploring the links between functional traits, life history traits, and fitness across species and environments would contribute to an integrated understanding of functional and life history strategies (Adler et al., 2014; Kelly et al., 2021).

A limitation of our approach is that the fitness estimates are difficult to externally validate. In one sense, λ is an integrated measure of demographic performance that is mathematically derived from size-specific demographic rates, so the λ estimates are as valid as the demographic rate estimates themselves (Caswell, 2001). In another sense, λ is the growth rate of a population at its equilibrium size structure in a stable environment, which are fairly strong assumptions. Our framework could be used to project short-term population growth rates and changes in population size structure, which could be validated using population time series data. The general ability of a model to reproduce realistic community dynamics could also be validated by comparing the composition and structure of simulated vs. observed communities.

However, because real communities are structured by historical processes not captured in our modeling framework (e.g., dispersal, drift, selection in past environments), a mismatch would not necessarily invalidate a model’s ability to estimate population fitness in current or projected future environments.

## Conclusions

Here we have proposed a framework for estimating the effects of multidimensional phenotypes on fitness across species. By integrating the effects of traits on demographic performance across species and over the life cycle into a single population model, this approach allows estimation of the net effects of traits on population fitness, revealing the contours of fitness landscapes and how they vary across environmental gradients. Our approach is flexible and can be applied in any system given the availability of trait and demographic data, which are becoming more widely available due to the proliferation of global databases (e.g., Kattge et al., 2020; Salguero-Gómez et al., 2015), providing a promising pathway to achieve the long-held goal of making community ecology more general, mechanistic, and predictive.

## Acknowledgements

We thank Stephen Ellner for modelling feedback and advice. This study was funded by the National Science Foundation EPSCOR Grant #2019528.

## Data availability

All data and code necessary to recreate the analyses presented in this manuscript are available on GitHub (http://github.com/andrewsiefert/treescapes).

## Author contributions

AS and DCL conceived the ideas and designed methodology; AS assembled and analyzed the data; AS and DCL wrote the manuscript.

