## Supporting Information for "Estimating the net effect of functional traits on fitness across species and environments"

Andrew Siefert* and Daniel C. Laughlin

Department of Botany, University of Wyoming, Laramie, WY 82071

**Supplementary methods**

**Data**

We fit demographic models using data from the US Forest Service Forest Inventory and Analysis (FIA; https://apps.fs.usda.gov/fia/datamart/datamart.html). We included plots on forest land with stand age of at least 20 years in the Northeastern Mixed Forest, Eastern Broadleaf Forest, and Southeastern Mixed Forest ecoregions and “rolling uplands” physiographic class. We excluded plots with evidence of human disturbance. FIA plots are repeatedly censused on cycles whose timing varies by state. For each state, we used the census cycle that included the year 2014, ensuring that data for all states came from a completed census cycle conducted using nationally standardized protocols (established in 1999). We excluded plots with a census interval shorter than 4 years. This left 12,752 plots in 20 states. Each FIA plot consists of four 7.32-m radius subplots in which all trees ≥12.7 cm diameter at breast height (dbh; “canopy trees” hereafter) are censused. Each subplot contains a 2.07-m radius microplot in which saplings of diameter ≥2.54 cm and <12.7 cm are censused. We excluded trees that were killed or damaged by human activities during the census interval. Canopy trees and saplings are tagged and tracked between censuses, allowing individual survival and growth to be measured over the census interval.

We extracted data on wood density, specific leaf area, and maximum height for tree species in FIA plots from the TRY plant trait database (Kattge *et al.* 2020). We joined species trait data with FIA data to create datasets for demographic modeling. Wood density and maximum height were log transformed prior to analysis.

**Demographic models**

***Survival***

The relationship between size and survival was not well described across the entire range of tree sizes by several functional forms we tried, so we built separate models for saplings (dbh < 12.7 cm) and canopy trees (dbh ≥ 12.7 cm). For both saplings and canopy trees, we modeled survival as:

$$\mathrm{survival}_{i} \sim\mathrm{Bernoulli}(p_{i})$$

$p_{i}= \left( \frac{S_{i}}{1+S_{i}} \right)^{{RI}_{i}}$	(1)

where *p_i_* is the expected survival probability of individual *i* and *RI_i_* is the census interval for individual *i*, in years. *S_i_* determines the expected 1-year survival probability of individual *i*. Building on models of Canham & Murphy (2016, 2017), we modeled survival as a multiplicative function of terms representing the potential survival rate of species *s* in plot *p* when all other predictors are at their average values (*PS_sp_*) and effects of size, crowding, climate, and traits:

$S_{isp}={PS}_{sp}\times{size}_{is}\times{crowding}_{is}\times{climate}_{sp}\times{trait}_{sp}$	(2)

Potential survival was modeled as:

${PS}_{sp}=exp(\alpha_{\mathrm{PS}}+\gamma_{\mathrm{PS}_{s}}+\gamma_{\mathrm{PS}_{p}})$	(3)

where *α*_PS_ determine the average potential survival, $\gamma_{\mathrm{PS}_{s}}$ is a random effect for species *s*, and $\gamma_{\mathrm{PS}_{p}}$ is a random effect for plot *p*.

For canopy tree survival, we modeled the ontogenetic effect of tree size as:

${size}_{is}=exp\left( {\beta_{size1}}_{s}{\log(\mathrm{dbh}}_{i})+\beta_{{size2}_{s}}\mathrm{dbh}_{i} \right)$	(4)

where *size_is_* is the ontogenetic effect of size on the survival of individual *i* of species *s*, dbh*_i_* is the diameter of individual *i* at the beginning of the census interval, and ${\beta_{size1}}_{s}$ and $\beta_{{size2}_{s}}$ are coefficients defining the shape of the size curve for species *s*. The function is hump-shaped (survival probability increases with size up to an optimum size, beyond which survival decreases) when *β*_size1_ is positive and *β*_size2_ is negative.

For saplings, we assumed that survival increased monotonically with size:

${size}_{is}={\mathrm{dbh}_{i}}^{{\beta_{size1}}_{s}}$	(5)

The parameters of the size functions were modeled as functions of traits, allowing for trait-by-size interactions, with species random effects:

${\beta_{size1}}_{s}=\beta_{size1}+\boldsymbol{\delta}_{size1}\mathbf{trait}_{s}+\gamma_{{size1}_{s}}$	(6)

${\beta_{size2}}_{s}=\beta_{size2}+\boldsymbol{\delta}_{size2}\mathbf{trait}_{s}+\gamma_{{size2}_{s}}$	(7)

where *β*_size1_ and *β*_size1_ are average size effects, ***δ***_size1_ and ***δ***_size2_ are vectors of trait-by-size interaction coefficients, **trait***_s_* is the vector of trait values for species *s*, and $\gamma_{{size1}_{s}}$ and $\gamma_{{size2}_{s}}$ are species random effects.

The effect of crowding was modeled as a power law function of neighbor basal area:

${crowding}_{is}={\mathrm{BA}_{i}}^{{\beta_{\mathrm{crowd}}}_{s}}$	(8)

where *crowding_is_* is the effect of crowding on the survival of individual *i* of species *s*, BA*_i_* is the total basal area of canopy trees in the same plot as individual *i*, and ${\beta_{\mathrm{crowd}}}_{s}$ defines the response to crowding of species *s*, modeled as:

${\beta_{\mathrm{crowd}}}_{s}=\beta_{\mathrm{crowd}}+\gamma_{\mathrm{crowd}_{s}}$	(9)

where *β*_crowd_ is the average crowding response and $\gamma_{\mathrm{crowd}_{s}}$ is the random effect for species *s*.

The effect of mean annual temperature was modeled as:

${climate}_{sp}=\exp\left( \beta_{\mathrm{clim}_{s}}\mathrm{MAT}_{p} \right)$	(10)

where MAT*_p_* is the mean annual temperature of plot *p* and $\beta_{\mathrm{clim}_{s}}$ defines the growth response to temperature of species *s*, modeled as:

$\beta_{\mathrm{clim}_{s}}=\beta_{\mathrm{clim}}+\gamma_{\mathrm{clim}_{s}}$	(11)

where *β*_clim_ is the average temperature response and $\gamma_{\mathrm{clim}_{s}}$ is the random effect for species *s*.

The effect of traits was modeled as a Gaussian (exponentiated quadratic) surface (Lande 1980):

$trait_{sp}=\exp\left( \boldsymbol{\beta}_{\mathrm{dir}_{p}}^{\boldsymbol{\top}}\mathbf{trait}_{s}+\mathbf{trait}_{s}^{\boldsymbol{\top}}\beta_{\mathrm{nonlin}_{p}}\mathbf{trait}_{s}) \right.$	(12)

where *trait_sp_* is the effect of traits on the survival probability of species *s* in plot *p*, $\boldsymbol{\beta}_{\mathrm{dir}_{p}}$ is a vector of directional (linear) performance gradients in plot *p*, and $\boldsymbol{\beta}_{\mathrm{nonlin}_{p}}$ is a matrix of nonlinear performance gradients in plot *p* (Arnold 2003). The diagonal elements of $\boldsymbol{\beta}_{\mathrm{nonlin}}$ measure the strength of stabilizing (if *β* is negative) or disruptive (if *β* is positive) selection for each trait, and the off-diagonal elements measure the strength of correlational selection between trait pairs(Arnold 2003). Positive correlational selection means that performance is maximized by having either high or low values of both traits. Negative correlational selection means that performance is maximized by having a high value of one trait and low value of the other trait. This function can produce performance landscapes of various shapes, including (when viewed in 2 dimensions) a peak, a saddle, a ridge, or a slope.

To allow performances surfaces to vary across the temperature gradient, the performance surface parameters (elements of $\boldsymbol{\beta}_{\mathrm{dir}}$ and $\boldsymbol{\beta}_{\mathrm{nonlin}}$) were modeled as linear functions of mean annual temperature:

$\beta_{\mathrm{dir}_{t,p}}=\beta_{\mathrm{dir}_{t}}+\delta_{\mathrm{dir}_{t}}\mathrm{MAT}_{p}$	(13)

$\beta_{\mathrm{nonlin}_{tu,p}}=\beta_{\mathrm{nonlin}_{tu}}+\delta_{\mathrm{nonlin}_{tu}}\mathrm{MAT}_{p}$	(14)

where $\beta_{\mathrm{dir}_{t,p}}$ is the directional selection coefficient for trait *t* in plot *p*, $\beta_{\mathrm{nonlin}_{tup}}$ is the stabilizing/disruptive selection coefficient for trait *t* (if *t* = *u*) or correlational selection coefficient for traits *t* and *u* (if *t* ≠ *u*) in plot *p*. $\beta_{\mathrm{dir}_{t}}$ and $\beta_{\mathrm{nonlin}_{tu}}$ are the performance gradients in a plot with average temperature, and $\delta_{\mathrm{dir}_{t}}$ and $\delta_{\mathrm{nonlin}_{tu}}$ describe how the trait effects change along the temperature gradient (i.e., trait-by-environment interactions).

***Growth***

Because growth rates could be zero or positive (we excluded negative growth values as we assume they resulted from damage), we modeled growth as a hurdle Gamma distribution:

$\Pr\left( \mathrm{growth}_{i} \right)= \left\{ \begin{aligned} \theta_{i} \mathrm{if} &\mathrm{growth}_{i}=0 \\ \left( 1- \theta_{i} \right)*\mathrm{Gamma}\left( G_{i}\beta, \beta\right) {\mathrm{if} \mathrm{growth}}_{i}>0 \end{aligned} \right.$	(15)

where *θ_i_* is the probability of zero growth for individual *i*, *G_i_* is the expected annual growth rate of individual *i*, and *β* is the rate parameter of the Gamma distribution. We assumed that both parts of the hurdle model (probability of zero growth and probabilities of positive growth values) are determined by the same processes. Therefore, instead of modeling them independently as commonly done in hurdle models, we modeled *θ* as a function of *G*:

$\theta_{i}= \frac{1}{1 + e^{-(z-G_{i})}}$	(16)

where *z* is an estimated parameter. In this formulation, individuals with high expected growth (*G_i_*) have a low probability of zero growth.

We modeled expected growth as a multiplicative function of terms representing the potential growth rate when all predictors are at their average values (*PG_sp_*) and effects of size, crowding, climate, and traits:

$G_{isp}={PG}_{sp}\times{size}_{is}\times{crowding}_{is}\times{climate}_{sp}\times{trait}_{sp}$	(17)

Potential growth was modeled as:

${PG}_{sp}=\exp(\alpha_{\mathrm{PG}}+\gamma_{\mathrm{PG}_{s}}+\gamma_{\mathrm{PG}_{p}})$	(18)

where *α*_PG_ is the overall average, $\gamma_{\mathrm{PG}_{s}}$ is a random effect for species *s*, and $\gamma_{\mathrm{PG}_{p}}$ is a random effect for plot *p*.

The terms for effects of size, crowding, climate, and traits on growth rate had the same form as the corresponding terms in the canopy survival model described above.

***Recruitment***

We modeled plot-level recruitment as a negative binomial distribution:

$\mathrm{recruitment}_{sp} = \mathrm{NegBinomial}\left( {recr}_{sp}*{RI}_{p}, \phi\right)$	(19)

where *recr_sp_* is the expected annual recruitment of species *s* in plot *p*, *RI_p_* is the recensus interval of plot *p* in years, and *ϕ* is an estimated parameter that controls overdispersion.

The expected annual recruitment rate of species *s* in plot *p* was determined by summing the expected annual per-capita recruitment rates of censused trees in the plot:

${recr}_{sp} = \sum_{i = 1}^{n} R_{isp}$	(20)

where *R_ksp_* is the annual production of new recruits by tree *i* of species *s* in plot *p*. Similar to survival and growth, we modeled expected annual per-capita recruitment as a multiplicative function of terms representing the potential recruitment rate and effects of size, crowding, climate, and traits:

$R_{isp}={PR}_{sp}\times{size}_{is}\times{crowding}_{is}\times{climate}_{ps}\times{trait}_{sp}$	(21)

Potential recruitment was modeled as:

${PR}_{sp}=\exp(\alpha_{\mathrm{PR}}+ \gamma_{\mathrm{PR}_{s}}+\gamma_{\mathrm{PR}_{p}})$	(22)

where *α*_PR_ is the overall average, $\gamma_{\mathrm{PR}_{s}}$ is a random effect for species *s*, and $\gamma_{\mathrm{PR}_{p}}$ is a random effect for plot *p*.

The size effect was modeled as the product of a term representing the effect of size on the probability of being reproductive and a term representing the effect of size on per-capita production of recruits, conditional on being reproductive (Ribbens *et al.* 1994; Clark *et al.* 2004):

${size}_{is}=\mathrm{logit}^{-1}\left( \beta_{\mathrm{repro}_{s}}*\left( \log\left( \mathrm{dia}_{i} \right)-D_{s} \right) \right) \times{\mathrm{dia}_{i}}^{{\beta_{recr}}_{s}}$	(23)

where *size_is_* is the ontogenetic effect of size on production of new recruits by individual *i* of species *s*, dia*_i_* is the diameter of individual *i* (in cm/30), $\beta_{\mathrm{repro}_{s}}$ is the slope of the effect of size on the probability of being reproductive for species *s*, *D_s_* is the log-diameter at which individuals of species *s* have a 50% chance of being reproductive, and ${\beta_{recr}}_{s}$ defines the effect of size on per-capita production of new recruits for species *s*.

The parameters of this size function varied among species as a function of species’ trait values, allowing trait-by-size interactions:

$D_{s}=D+\boldsymbol{\delta}_{D}\mathbf{trait}_{s}+\gamma_{D_{s}}$	(24)

${\beta_{\mathrm{repro}}}_{s}= \exp\left( \beta_{\mathrm{repro}}+\boldsymbol{\delta}_{\mathrm{repro}}\mathbf{trait}_{s}+\gamma_{\mathrm{repro}_{s}} \right)$	(25)

${\beta_{\mathrm{recr}}}_{s}=\exp\left( \beta_{\mathrm{recr}}+\boldsymbol{\delta}_{\mathrm{recr}}\mathbf{trait}_{s}+\gamma_{\mathrm{recr}_{s}} \right)$	(26)

where *D*, *β*_repro_, and *β*_recr_ are parameters of the size function for species with average trait values, the ***δ***’s are vectors of trait effects on those parameters (trait-by-size interactions), and the *γ*’s are species random effects.

The parameters describing the effect of size on the probability of being reproductive were estimated jointly using plot-level FIA recruitment data and individual-level size and reproductive status data for trees in the MASTIF data set (Clark *et al.* 2021). Reproductive status of MASTIF trees was modeled as:

$\mathrm{repro}_{i} \sim\mathrm{Bernoulli}(pr_{i})$	(27)

${pr}_{jqsp}=\mathrm{logit}^{-1}\left( \beta_{\mathrm{repro}_{s}}*\left( \log\left( \mathrm{dia}_{i} \right)-D_{s} \right)+\gamma_{\mathrm{site}_{p}}+\gamma_{\mathrm{tree}_{q}} \right)$	(28)

where *pr_jqsp_* is the probability of being reproductive for observation *j* of tree *q* (some trees were observed in multiple years) of species *s* in site *p*, and $\gamma_{\mathrm{site}_{p}}$ and $\gamma_{\mathrm{tree}_{q}}$ are random effects for site and tree, respectively. The site and tree random effects are assumed to be drawn from normal distributions with mean of zero and standard deviation *τ*_site_ and *τ*_tree_, respectively.

The effect of crowding on per-capita recruitment was modeled as a function of canopy and sapling neighbor basal area:

${crowding}_{is}={\mathrm{BA}_{\mathrm{can}_{i}}}^{{\beta_{\mathrm{can}}}_{s}} \times{\mathrm{BA}_{\mathrm{sap}_{i}}}^{{\beta_{\mathrm{sap}}}_{s}}$	(29)

were $\mathrm{BA}_{\mathrm{can}_{i}}$ and $\mathrm{BA}_{\mathrm{sap}_{i}}$ are the total basal areas of canopy trees and saplings, respectively, in the same plot as individual *i*, and ${\beta_{\mathrm{can}}}_{s}$ and ${\beta_{\mathrm{sap}}}_{s}$ define the response of species *s* to crowding by canopy trees and saplings, respectively. Species’ responses to crowding were modeled as:

${\beta_{\mathrm{can}}}_{s}=\beta_{\mathrm{can}}+\gamma_{\mathrm{can}_{s}}$	(30)

${\beta_{\mathrm{sap}}}_{s}=\beta_{\mathrm{sap}}+\gamma_{\mathrm{sap}_{s}}$	(31)

where *β*_can_ and *β*_sap_ are average responses to crowding by canopy trees and saplings, respectively, and the *γ*’s are species random effects.

The terms for climate and trait effects on recruitment had the same form as the corresponding terms in the survival and growth models.

For each model, plot-level random effects were assumed to be drawn from a normal distribution with mean of zero and standard deviation *τ*_plot_. Species-level random effects were assumed to be drawn from a multivariate normal distribution with means of zero and covariance matrix **Σ**:

$\text{Σ}=\mathrm{diag}\left( \boldsymbol{\tau}_{\mathrm{species}} \right)\boldsymbol{L}\boldsymbol{L}^{\boldsymbol{\top}}\mathrm{diag}\left( \boldsymbol{\tau}_{\mathrm{species}} \right)$	(32)

where ***τ***_species_ is a vector of standard deviations of species random effects and ***L*** is the Cholesky factor of the correlation matrix of species random effects.

***Model fitting***

For each model, we assigned weakly informative priors that constrained predicted demographic rates to biologically realistic values, as assessed by simulating data from the prior distributions. Given the amount of data used to fit the models, the priors had negligible effect on the posterior parameter estimates but aided in model convergence. We obtained posterior samples using Hamiltonian Monte Carlo implemented in Stan(Team n.d.). For each model, we ran 4 chains with 2,000 iterations, 1,000 of which were warmup. We assessed model convergence using the $\hat{R}$statistic (Vehtari *et al.* 2021; values for all parameters were ≤1.02).

**Fitness estimates**

We constructed IPM kernels for hypothetical populations with different combinations of trait and environmental values using the demographic rate models described above, with crowding effects held constant at their average value and species and plot random effects set to zero. For the survival kernel, *s(z)*, we used the sapling survival model for individuals with size *z* < 12.7 cm and the canopy survival model for individuals with size ≥12.7 cm. We constructed the recruitment kernel *F*(*z’,z*) as the product of the number of new recruits produced annually by an individual of size *z*, estimated using our recruitment model, and the size distribution of new recruits, which we modeled as a Gamma distribution fit to all recruits in the data set. It is possible that traits, environment, or initial size could affect recruit size, but we chose not to model these effects since simulations showed that the recruit size distribution had a negligible effect on projected population growth rates.

We implemented the IPMs using numerical integration (midpoint rule; Ellner *et al.* 2016). This method discretizes the kernel by dividing the size range into a large number of evenly-sized bins and evaluating the kernel at the midpoint of each bin, effectively generating a large matrix population model. We used a size range from 2.5 to 100 cm and 1000 bins, resulting in a 1000 x 1000 transition matrix. A matrix this large is necessary due to the large size range and small growth increments of trees. We confirmed that 1000 bins were enough to produce stable numerical results. To avoid “eviction” of individuals with predicted sizes greater than 100 cm (Ellner *et al.* 2016), we treated the largest size bin as a discrete size class that included all individuals with predicted sizes greater than 100 cm. The growth and recruit size distributions are probability distributions and should thus integrate to one (or sum to one in the discretized version) for any initial size *z_i_*, but discretizing the kernels resulted in this not being true in all cases. To correct this, for each initial size class *z_i_* we divided the probabilities of growing into (or new recruits being within) each size class *z_j_* by the sum of those probabilities, thus ensuring that the growth and recruit size probabilities for any initial size *z_i_* summed to one.

The population growth rate (λ), our estimate of fitness, was extracted as the dominant eigenvalue of the discretized kernel (Ellner *et al.* 2016). To estimate fitness landscapes, we constructed kernels and extracted population growth rates for a grid of trait values covering the observed trait space at each of three mean annual temperatures (5, 10, and 15°C). To quantify expected λ values, we constructed kernels using posterior mean parameter values from the demographic rate models. To estimate uncertainty in λ, we constructed kernels using 800 samples from the posteriors of the demographic rate models and calculated 90% credible intervals.

**λ estimates for single species**

To confirm that the FIA data set captures variation in species’ performance in response to mean annual temperature, we estimated λ for four representative tree species across their temperature ranges in the FIA data. We chose species that had ample demographic data from FIA and reproductive status data from MASTIF and had clear northern (*Liquidambar styraciflua* and *Ulmus alata*) or southern (*Picea rubens* and *Betula alleghaniensis*) range limits within the study area. For each species, we fit demographic rate models similar to the multi-species models described above, except we included a quadratic term for mean annual temperature and omitted trait effects and species random effects. We constructed IPM kernels using draws from the posterior distributions of the demographic rate models for each species at 20 mean annual temperature values covering their respective ranges and extracted λ values as described above.

**Supplementary figures**

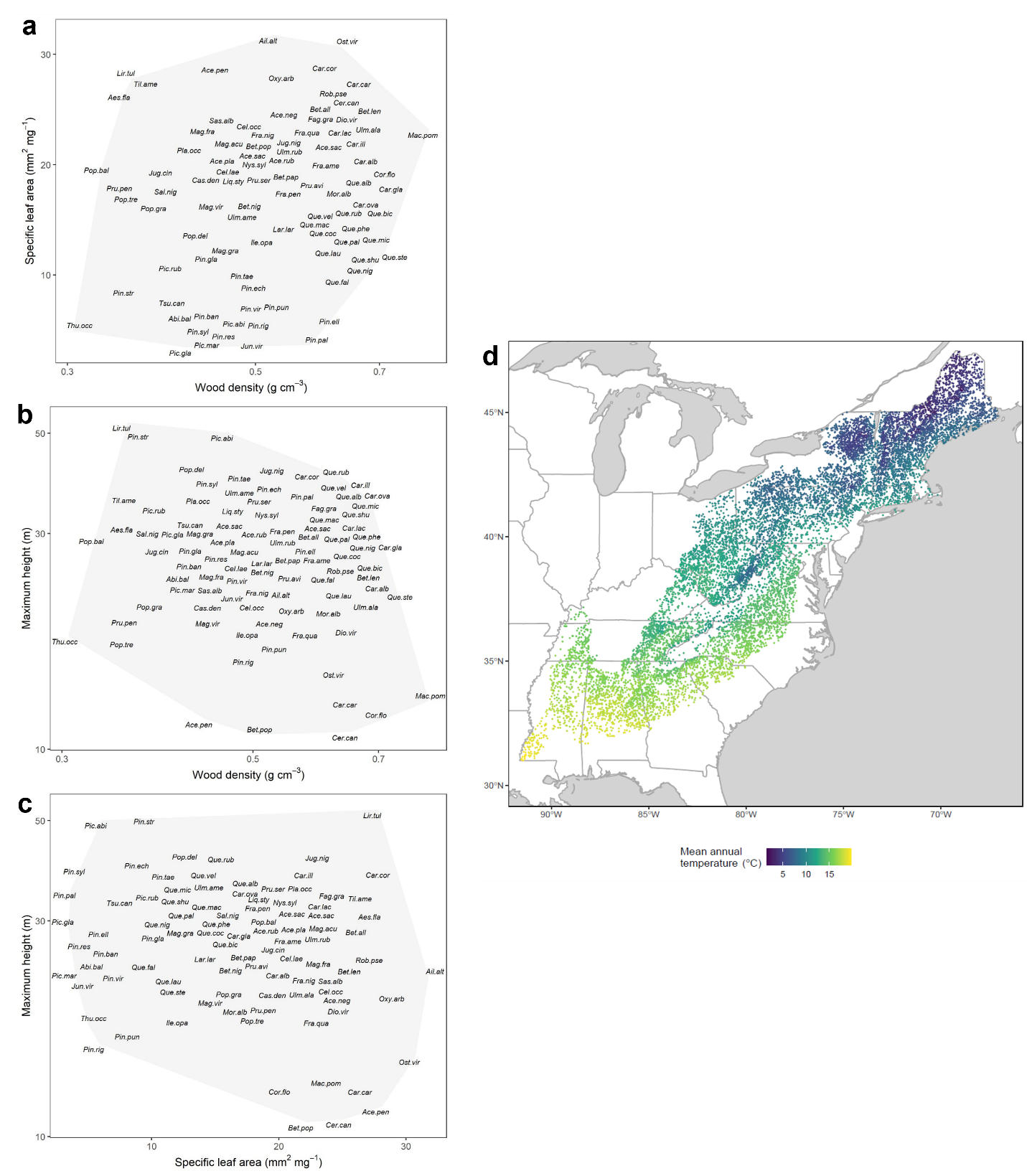

**Figure S1.** Maps of tree species in trait space and plots in geographic space. Trait space maps show locations of species in two-dimensional trait space defined by a) wood density and specific leaf area (SLA), b) wood density and maximum height, or c) SLA and maximum height. d) Map of US Forest Inventory and Analysis (FIA) plots included in the data set for modeling tree demographic rates.

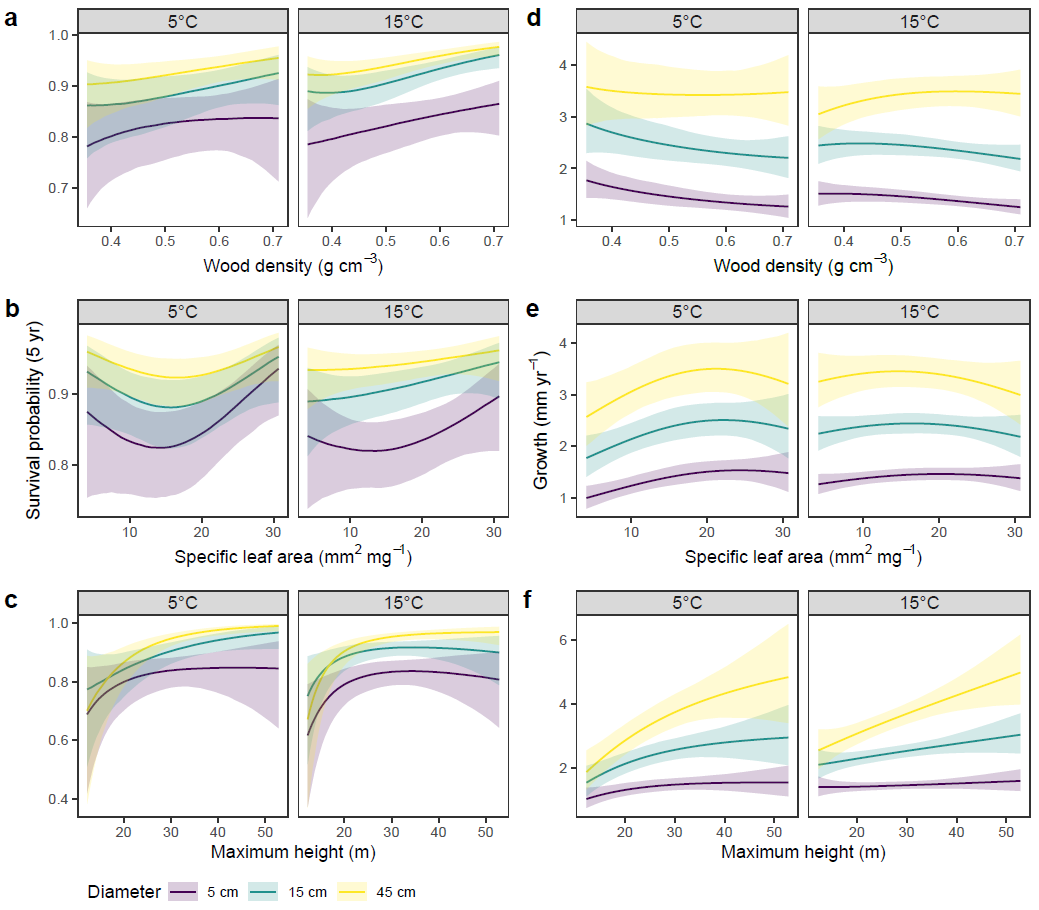

**Figure S2.** Functional trait effects on survival and growth with respect to tree diameter and mean annual temperature. Plots show conditional effects of wood density (a,d), specific leaf area (b,e), or maximum height (c,f) on 5-year survival probability (a-c) or diameter growth rate (d-f) at low (5°C) and high (15°C) mean annual temperature with other traits and crowding effects held constant at their average values. Trend lines show posterior expectations and shaded areas show 90% credible intervals.

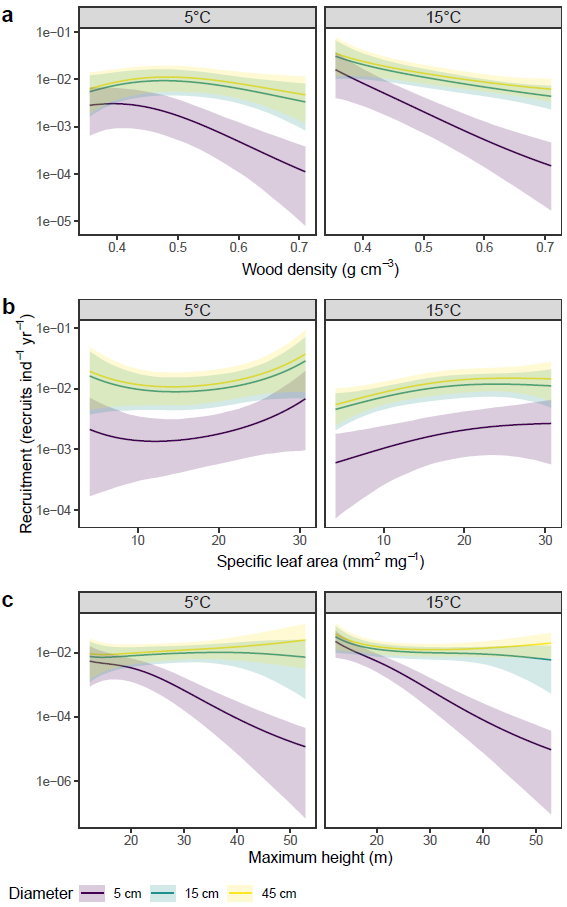

**Figure S3.** Trait effects on recruitment with respect to tree diameter and mean annual temperature. Plots show conditional effects of wood density (a), specific leaf area (b), or maximum height (c) on recruitment rate (number of recruits, i.e., saplings reaching 2.54 cm diameter, produced by an adult tree per year) at low (5°C) and high (15°C) mean annual temperature with the other traits and crowding effects held constant at their average values. Trend lines show posterior expectations and shaded areas show 90% credible intervals. The y-axis is log transformed to show trends more clearly.

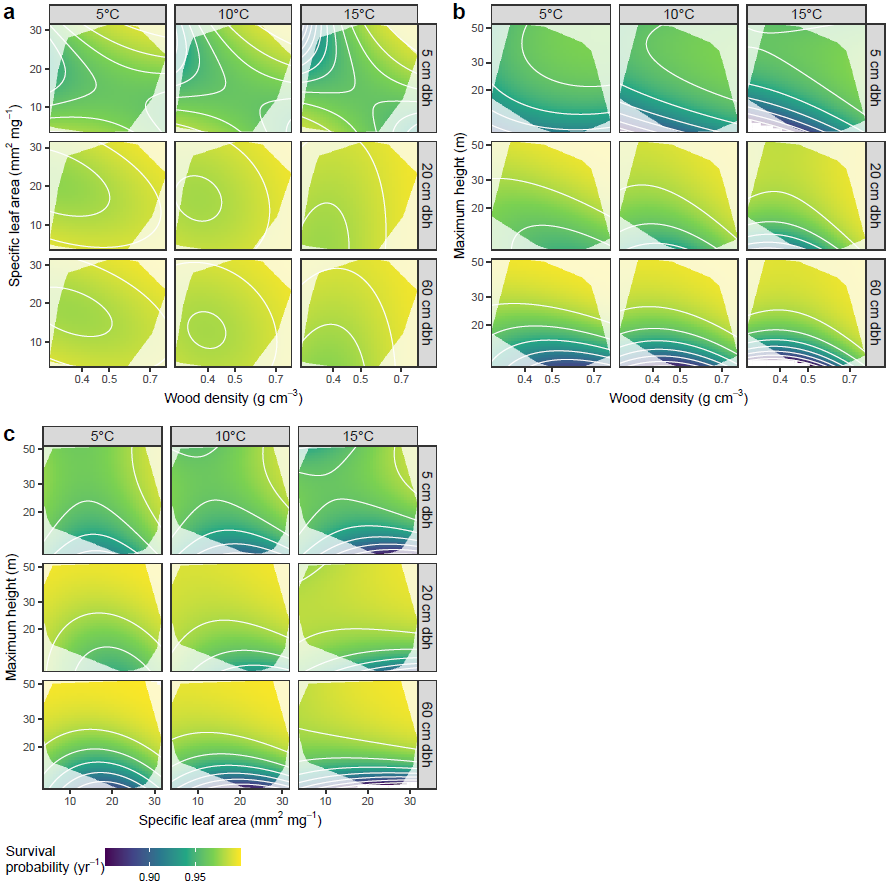

**Figure S4.** Survival performance landscapes with respect to tree diameter and mean annual temperature. Landscapes show expected 1-year survival probability for trees species with different combinations of trait values. Our survival model included three traits—wood density, specific leaf area (SLA), and maximum height—but for ease of visualization, landscapes are shown for two traits at a time (wood density and SLA, a; wood density and maximum height, b; SLA and maximum height, c) with the third trait held constant at its average value. Grayed areas show regions of trait space not occupied by tree species in our data set.

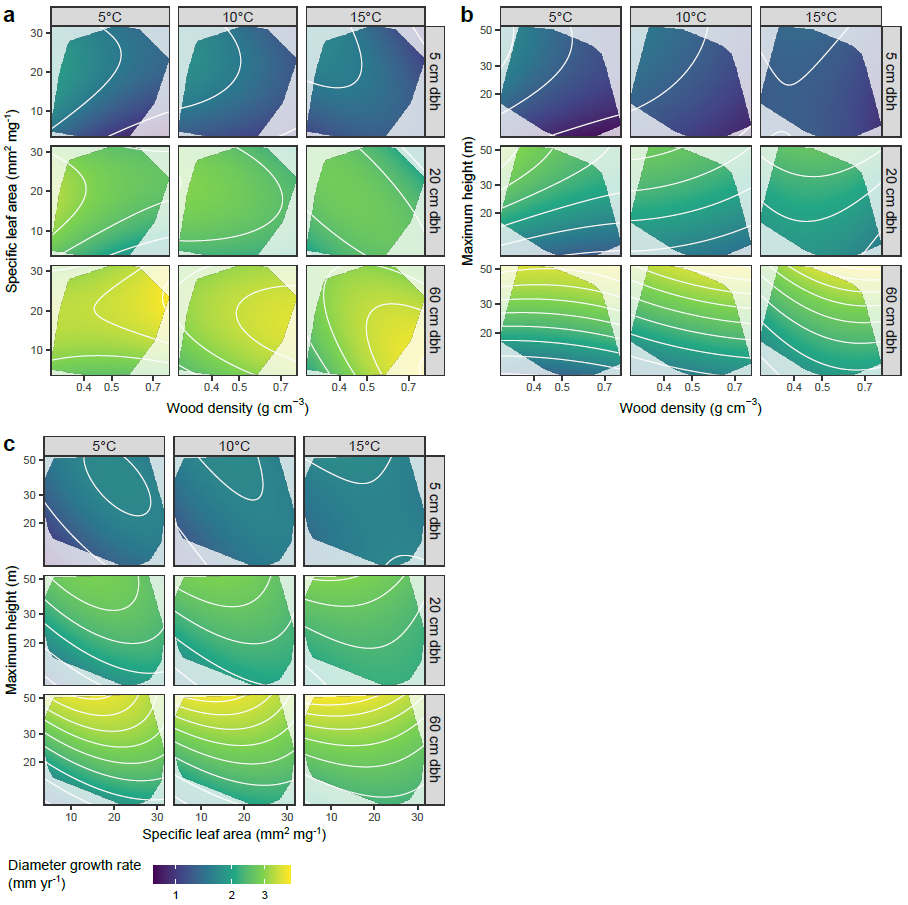

**Figure S5.** Growth performance landscapes by tree size and temperature. Landscapes show expected diameter growth rate for trees species with different combinations of trait values. Our growth model included three traits—wood density, specific leaf area (SLA), and maximum height—but for ease of visualization, landscapes are shown for two traits at a time (wood density and SLA, a; wood density and maximum height, b; SLA and maximum height, c) with the third trait held constant at its average value. Grayed areas show regions of trait space not occupied by tree species in our data set.

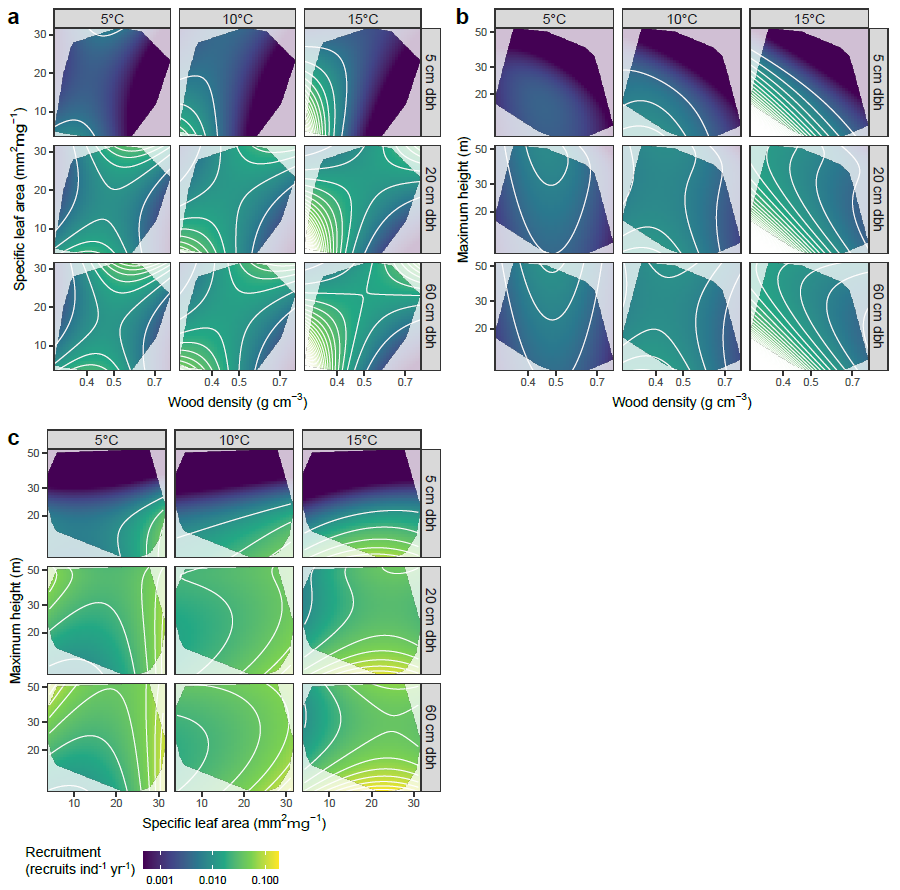

**Figure S6.** Recruitment performance landscapes by tree diameter and mean annual temperature. Landscapes show expected recruitment rate (number of recruits, i.e., saplings reaching 2.54 cm diameter, produced by an adult tree per year) for trees species with different combinations of trait values. Our growth model included three traits—wood density, specific leaf area (SLA), and maximum height—but for ease of visualization, landscapes are shown for two traits at a time (wood density and SLA, a; wood density and maximum height, b; SLA and maximum height, c) with the third trait held constant at its average value. Grayed areas show regions of trait space not occupied by tree species in our data set.

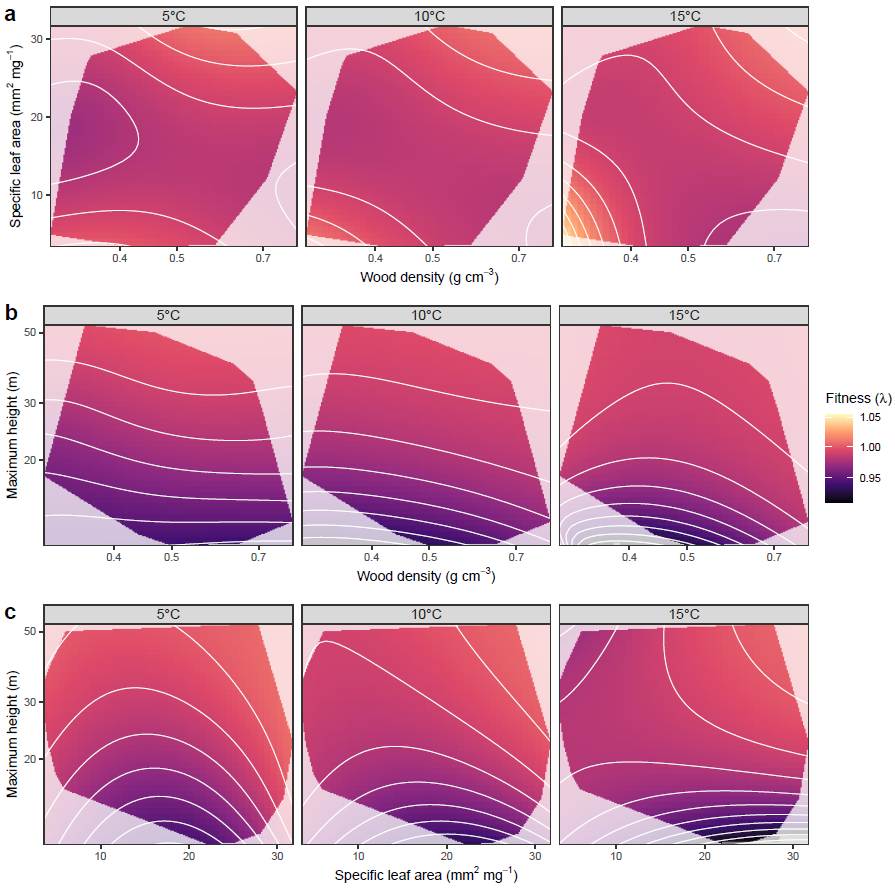

**Figure S7.** Fitness landscapes at low, medium, and high mean annual temperature. Landscapes show expected fitness (population growth rate, λ) for trees species with different trait combinations at low (5°C), medium (10°C), and high (15°C) mean annual temperatures and average neighbor density. Models included three traits—wood density, specific leaf area (SLA), and maximum height—but for ease of visualization, landscapes are shown for two traits at a time (wood density and SLA, a; wood density and maximum height, b; SLA and maximum height, c) with the third trait held constant at its average value. Grayed areas show regions of trait space not occupied by tree species in our data set.

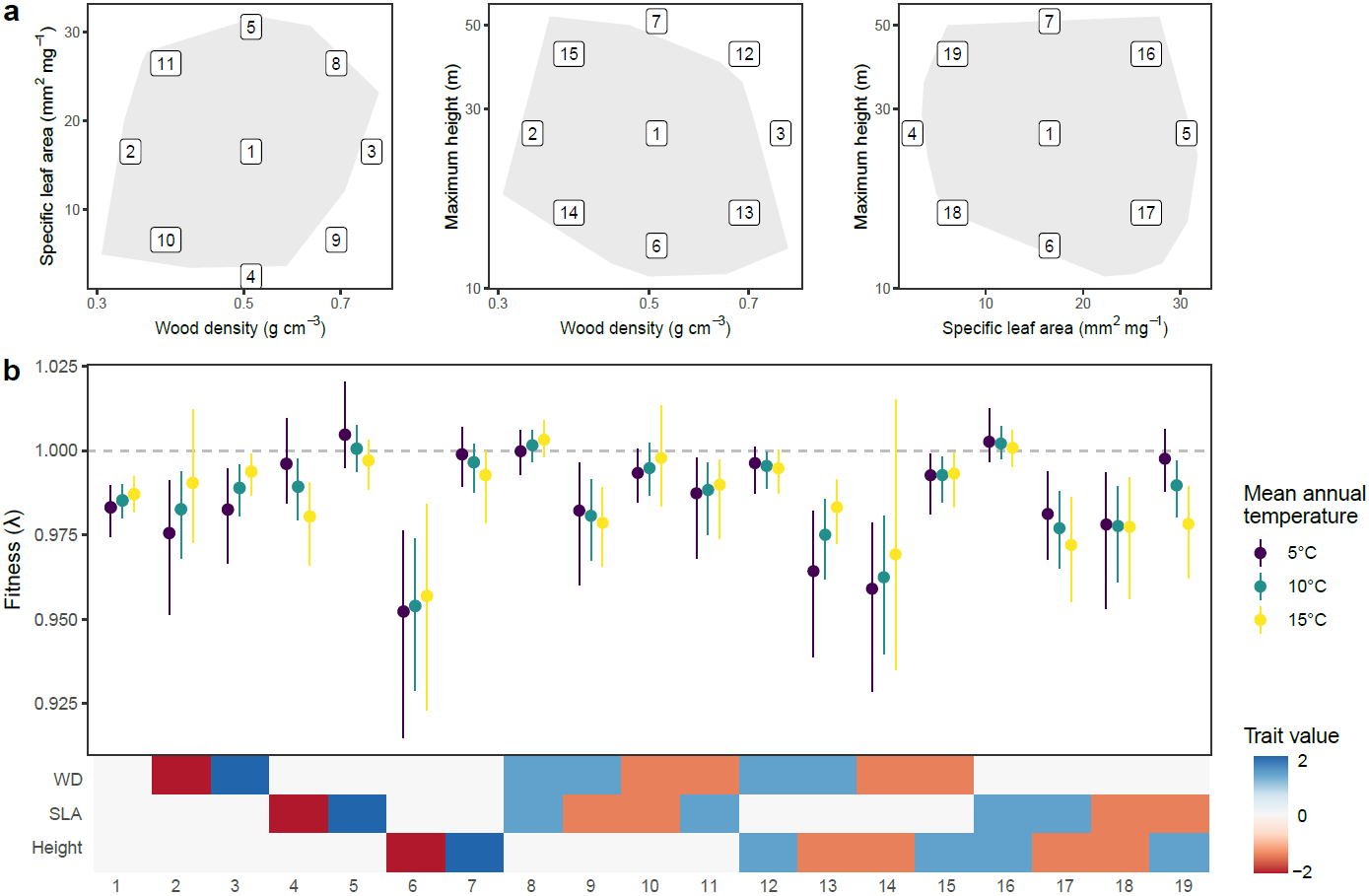

**Figure S8.** Uncertainty in fitness estimates for trait combinations in low, medium, and high mean annual temperature. To illustrate uncertainty in estimates of fitness (population growth rate, λ) from trait-based population projection models, we obtained fitness estimates for representative trait combinations (indicated by numbered points in trait space, panel a) at low (5°C), medium (10°C), and high (15°C) mean annual temperatures and average neighbor density. For each trait and temperature combination, we took 1000 samples from the posterior distributions of demographic rate (survival, growth, and recruitment) models. Then, for each set of sampled demographic parameters, we constructed a population projection model (IPM) and calculated λ, resulting in a posterior distribution for λ. Points and error bars in panel b show posterior means and 90% credible intervals, respectively, from this distribution. Numbered trait combinations shown on the *X*-axis of panel b correspond to numbered locations in trait space in panel a. Trait values, indicated by colored boxes below the *X*-axis of panel b, are scaled to mean of 0 and standard deviation of 1 to make relative values comparable across traits.

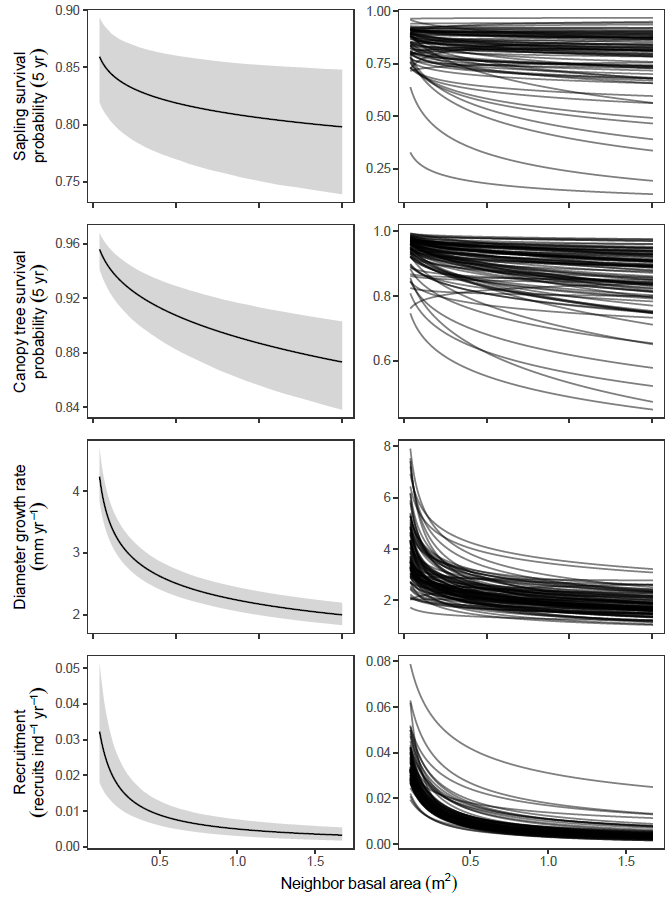

**Figure S9.** Effects of crowding on tree demographic rates. Plots show conditional mean effects (posterior mean expected values and 90% credible intervals) and species-specific effects (right panel) of crowding (measured as total basal area of neighboring canopy trees within a plot) on tree demographic rates, with other predictors (traits, tree diameter, temperature) held constant at their average values.

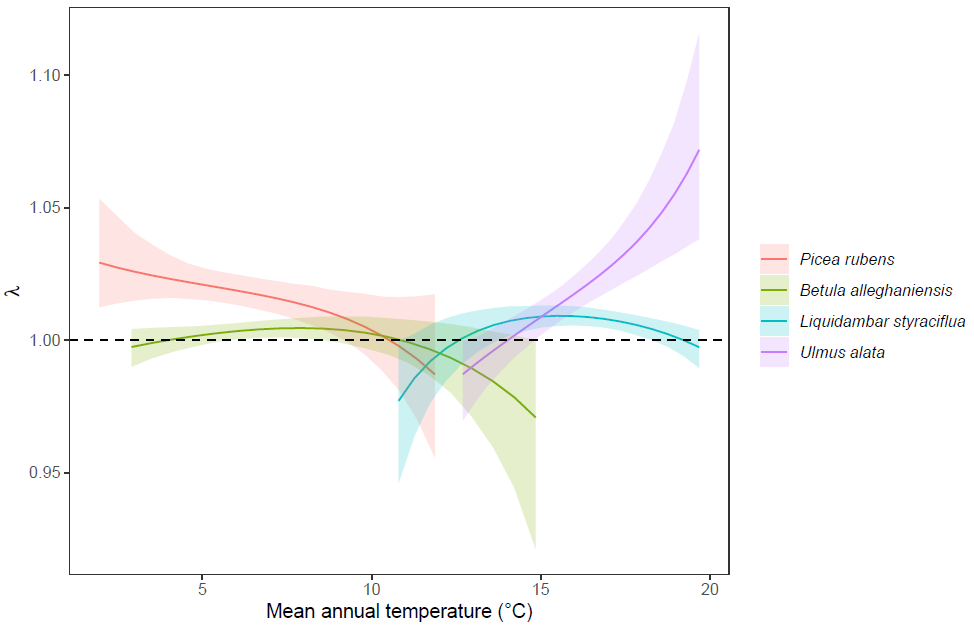

**Figure S10.** Population growth rate (λ) estimates for four representative tree species across a mean annual temperature gradient in the eastern United States.

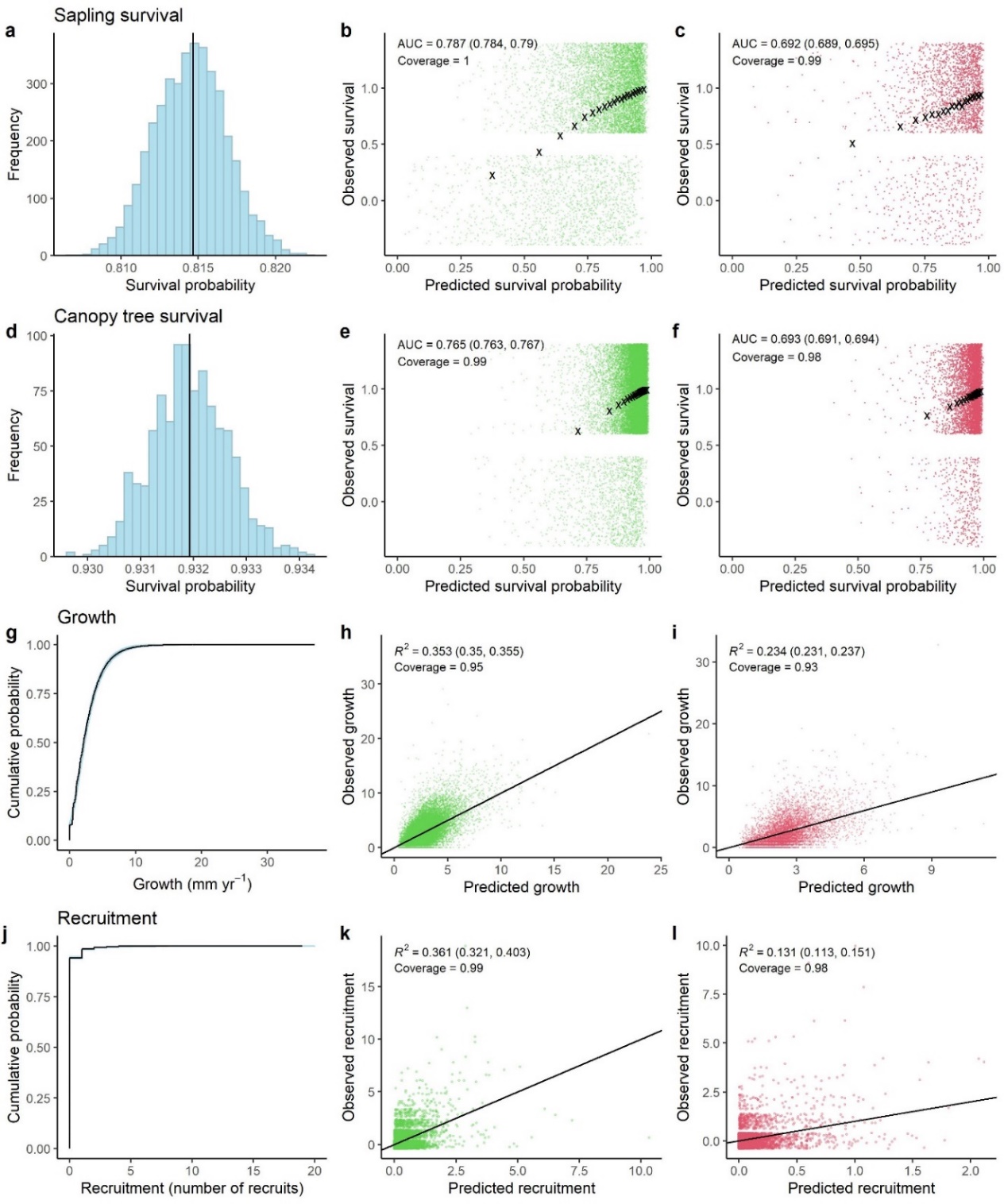

**Figure S11. Posterior checks and model validation for demographic models.** Posterior predictive checks for survival show the posterior distribution of mean survival across all trees in the data set. Posterior predictive checks for growth (g) and recruitment (j) show empirical cumulative distributions for observed data and posterior predictions. Panels b,e,h,j and c,f,i,l show the relationship between predicted (posterior means) and observed values for within-sample and out-of-sample data (20% of plots), respectively. Model fit for survival was assessed using area under the ROC curve (AUC; values displayed are posterior mean and 90% CI) and coverage (proportion of observations that fall within the 90% CI for predictions). Observed survival was jittered for clarity. Black x’s show mean predicted and observed survival for 20 equal-sized bins. Model fit for growth and recruitment were assessed using Bayesian R^2^ (values displayed are posterior mean and 90% CI) and coverage.

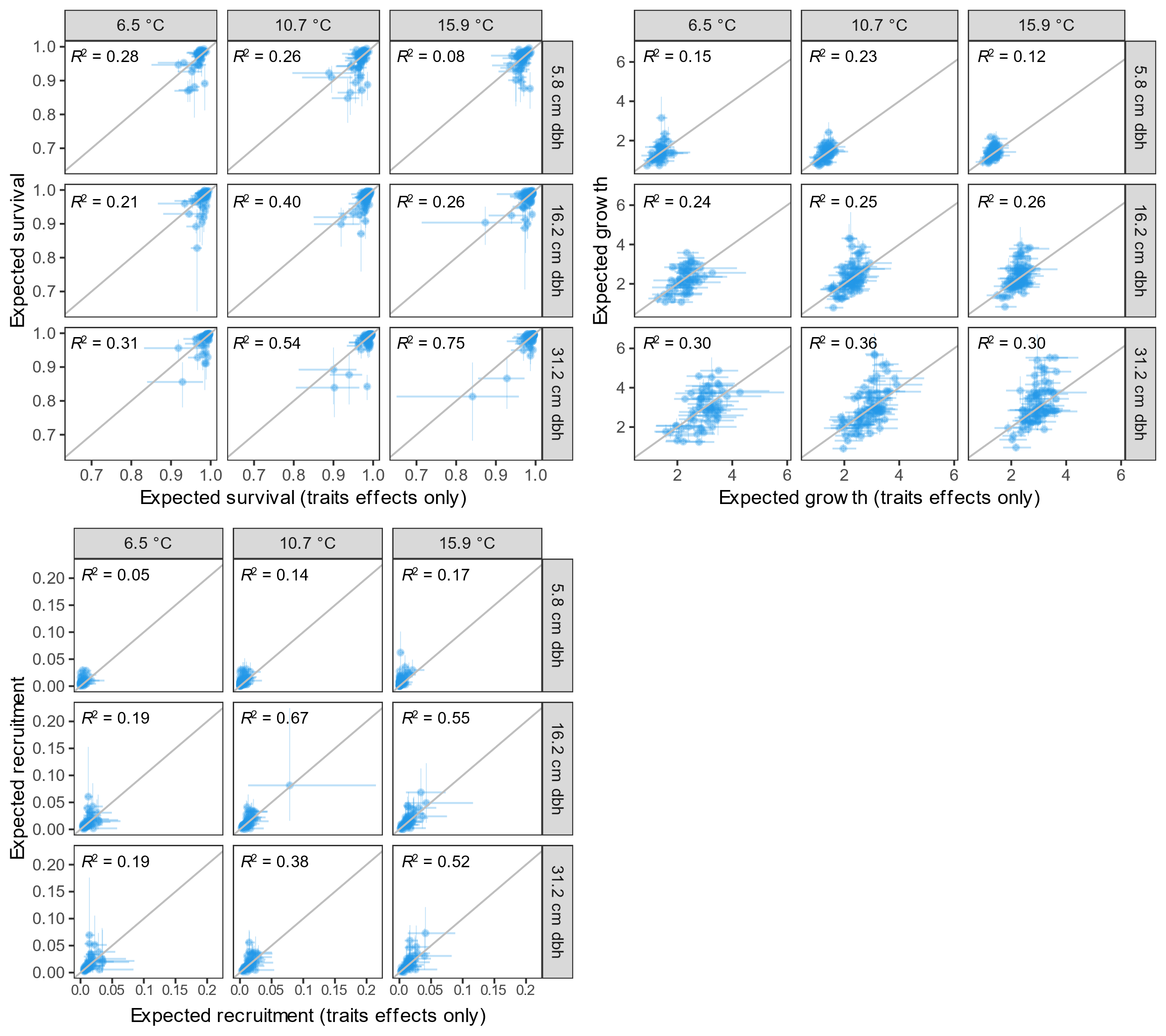

**Figure S12.** The proportion of variance in expected demographic rates explained by functional traits. To assess the degree to which traits explained variation in species’ demographic rates, we compared species’ expected annual survival probability (a), annual diameter growth rate (b), and annual per-capita recruitment rate (c) based on traits alone (i.e., all species random effects set to 0; x-axis) and traits plus species random effects (y-axis). To isolate the influence of traits, we binned the data based on size and mean annual temperature and calculated expected demographic rates for species within each bin with individual size and temperature held at their bin-specific averages and crowding held at its overall average. We calculated the proportion of variance in expected demographic rates explained by traits (*R*^2^) as the square of the Pearson correlation coefficient (*r*) between predictions based on traits alone and predictions based on traits plus species random effects within each bin. Points represent posterior means and error bars show 90% credible intervals.

**Table S1.** Descriptions of terms and variables in the sapling survival model. Terms are categorized as data (observed quantities), parameters (unobserved quantities that are directly sampled), and transformed parameters (unobserved quantities that are functions of parameters and data). Dimensions of vector and matrix terms are given in parentheses. Subscripts reference individual *i*, species *s*, plot *p*, trait *t* and *u*.

| **Term** | **Type** | **Description** |
| --- | --- | --- |
| $\mathrm{survival}_{i}$ | Data (response variable) | Observed survival |
| $p_{i}$ | Transformed parameter | Expected survival probability |
| ${RI}_{i}$ | Data | Recensus interval (years) |
| $S_{i}$ | Transformed parameter | Expected 1-year survival (transformed) |
| ${PS}_{sp}$ | Transformed parameter | Potential survival |
| $\alpha_{\mathrm{PS}}$ | Parameter | Average potential survival |
| $\gamma_{\mathrm{PS}_{s}}$ | Parameter | Random species effect on potential survival |
| $\gamma_{\mathrm{PS}_{p}}$ | Parameter | Random plot effect on potential survival |
| ${size}_{is}$ | Transformed parameter | Size term |
| $\mathrm{dbh}_{i}$ | Data | Diameter at breast height (cm) |
| ${\beta_{size1}}_{s}$ | Transformed parameter | log(size) coefficient |
| $\beta_{size1}$ | Parameter | Average log(size) coefficient |
| $\boldsymbol{\delta}_{\mathbf{size1}}$ (3) | Parameter | Trait effects on log(size) coefficient |
| $\gamma_{{size1}_{s}}$ | Parameter | Random species effect on log(size) coefficient |
| ${crowding}_{is}$ | Transformed parameter | Crowding term |
| $\mathrm{BA}_{i}$ | Data | Neighbor basal area (cm^2^) |
| ${\beta_{\mathrm{crowd}}}_{s}$ | Transformed parameter | Crowding response |
| $\beta_{\mathrm{crowd}}$ | Parameter | Average crowding response |
| $\gamma_{\mathrm{crowd}_{s}}$ | Parameter | Random species effect on crowding response |
| ${climate}_{sp}$ | Transformed parameter | Climate term |
| $\mathrm{MAT}_{p}$ | Data | Mean annual temperature (°C) |
| $\beta_{\mathrm{clim}_{s}}$ | Transformed parameter | Temperature response |
| $\beta_{\mathrm{clim}}$ | Parameter | Average temperature response |
| $\gamma_{\mathrm{clim}_{s}}$ | Parameter | Random species effect on temperature response |
| $trait_{sp}$ | Transformed parameter | Trait term (performance landscape) |
| $\mathbf{trait}_{s}$ (3) | Data | Trait vector (wood density, SLA, maximum height) |
| $\boldsymbol{\beta}_{\mathbf{dir}_{p}}$ (3) | Transformed parameter | Directional (linear) performance gradients |
| $\beta_{\mathrm{dir}_{t}}$ | Parameter | Average directional performance gradient |
| $\delta_{\mathrm{dir}_{t}}$ | Parameter | Temperature effect on directional performance gradient |
| $\boldsymbol{\beta}_{\mathbf{nonlin}_{p}}$ (3x3) | Transformed parameter | Nonlinear performance gradients |
| $\beta_{\mathrm{nonlin}_{tu}}$ | Parameter | Average nonlinear performance gradient |
| $\delta_{\mathrm{nonlin}_{tu}}$ | Parameter | Temperature effect on nonlinear performance gradient |
| $\tau_{\mathrm{plot}}$ | Parameter | Plot random effects standard deviation |
| $\boldsymbol{\Sigma}$ (4x4) | Transformed parameter | Species random effects covariance matrix |
| $\boldsymbol{\tau}_{\mathbf{species}}$ | Parameter | Species random effects standard deviations |
| ***L*** (4x4) | Parameter | Species random effects correlation matrix (Cholesky factor) |

**Table S2.** Descriptions of terms and variables in the canopy tree survival model. Terms are categorized as data (observed quantities), parameters (unobserved quantities that are directly sampled), and transformed parameters (unobserved quantities that are functions of parameters and data). Dimensions of vector and matrix terms are given in parentheses. Subscripts reference individual *i*, species *s*, plot *p*, trait *t* and *u*.

| **Term** | **Type** | **Description** |
| --- | --- | --- |
| $\mathrm{survival}_{i}$ | Data (response variable) | Observed survival |
| $p_{i}$ | Transformed parameter | Expected survival probability |
| ${RI}_{i}$ | Data | Recensus interval (years) |
| $S_{i}$ | Transformed parameter | Expected 1-year survival (transformed) |
| ${PS}_{sp}$ | Transformed parameter | Potential survival |
| $\alpha_{\mathrm{PS}}$ | Parameter | Average potential survival |
| $\gamma_{\mathrm{PS}_{s}}$ | Parameter | Random species effect on potential survival |
| $\gamma_{\mathrm{PS}_{p}}$ | Parameter | Random plot effect on potential survival |
| ${size}_{is}$ | Transformed parameter | Size term |
| $\mathrm{dbh}_{i}$ | Data | Diameter at breast height (cm) |
| ${\beta_{size1}}_{s}$ | Transformed parameter | log(size) coefficient |
| $\beta_{size1}$ | Parameter | Average log(size) coefficient |
| $\boldsymbol{\delta}_{\mathbf{size1}}$ (3) | Parameter | Trait effects on log(size) coefficient |
| $\gamma_{{size1}_{s}}$ | Parameter | Random species effect on log(size) coefficient |
| $\beta_{{size2}_{s}}$ | Transformed parameter | Size coefficient |
| $\beta_{size2}$ | Parameter | Average size coefficient |
| $\boldsymbol{\delta}_{\mathbf{size2}}$ (3) | Parameter | Trait effects on size coefficient |
| $\gamma_{{size2}_{s}}$ | Parameter | Random species effect on size coefficient |
| ${crowding}_{is}$ | Transformed parameter | Crowding term |
| $\mathrm{BA}_{i}$ | Data | Neighbor basal area (cm^2^) |
| ${\beta_{\mathrm{crowd}}}_{s}$ | Transformed parameter | Crowding response |
| $\beta_{\mathrm{crowd}}$ | Parameter | Average crowding response |
| $\gamma_{\mathrm{crowd}_{s}}$ | Parameter | Random species effect on crowding response |
| ${climate}_{sp}$ | Transformed parameter | Climate term |
| $\mathrm{MAT}_{p}$ | Data | Mean annual temperature (°C) |
| $\beta_{\mathrm{clim}_{s}}$ | Transformed parameter | Temperature response |
| $\beta_{\mathrm{clim}}$ | Parameter | Average temperature response |
| $\gamma_{\mathrm{clim}_{s}}$ | Parameter | Random species effect on temperature response |
| $trait_{sp}$ | Transformed parameter | Trait term (performance landscape) |
| $\mathbf{trait}_{s}$(3) | Data | Trait vector (wood density, SLA, maximum height) |
| $\boldsymbol{\beta}_{\mathbf{dir}_{p}}$ (3) | Transformed parameter | Directional (linear) performance gradients |
| $\beta_{\mathrm{dir}_{t}}$ | Parameter | Average directional performance gradient |
| $\delta_{\mathrm{dir}_{t}}$ | Parameter | Temperature effect on directional performance gradient |
| $\boldsymbol{\beta}_{\mathbf{nonlin}_{p}}$ (3x3) | Transformed parameter | Nonlinear performance gradients |
| $\beta_{\mathrm{nonlin}_{tu}}$ | Parameter | Average nonlinear performance gradient |
| $\delta_{\mathrm{nonlin}_{tu}}$ | Parameter | Temperature effect on nonlinear performance gradient |
| $\tau_{\mathrm{plot}}$ | Parameter | Plot random effects standard deviation |
| $\boldsymbol{\Sigma}$ (5x5) | Transformed parameter | Species random effects covariance matrix |
| $\boldsymbol{\tau}_{\mathbf{species}}$ (5) | Parameter | Species random effects standard deviations |
| ***L*** (5x5) | Parameter | Species random effects correlation matrix (Cholesky factor) |

**Table S3.** Descriptions of terms and variables in the growth model. Terms are categorized as data (observed quantities), parameters (unobserved quantities that are directly sampled), and transformed parameters (unobserved quantities that are functions of parameters and data). Dimensions of vector and matrix terms are given in parentheses. Subscripts reference individual *i*, species *s*, plot *p*, trait *t* and *u*.

| **Term** | **Type** | **Description** |
| --- | --- | --- |
| $\mathrm{growth}_{i}$ | Data (response variable) | Diameter growth rate (mm yr^-1^) |
| $G_{i}$ | Transformed parameter | Expected annual growth rate |
| $\theta_{i}$ | Transformed parameter | Probability of zero growth |
| $\beta$ | Parameter | Rate parameter of Gamma distribution |
| ${PG}_{sp}$ | Transformed parameter | Potential growth rate |
| $\alpha_{\mathrm{PG}}$ | Parameter | Average potential growth rate |
| $\gamma_{\mathrm{PG}_{s}}$ | Parameter | Random species effect on potential growth rate |
| $\gamma_{\mathrm{PG}_{p}}$ | Parameter | Random plot effect on potential growth rate |
| ${size}_{is}$ | Transformed parameter | Size term |
| $\mathrm{dbh}_{i}$ | Data | Diameter at breast height (cm) |
| ${\beta_{size1}}_{s}$ | Transformed parameter | log(size) coefficient |
| $\beta_{size1}$ | Parameter | Average log(size) coefficient |
| $\boldsymbol{\delta}_{\mathbf{size1}}$ (3) | Parameter | Trait effects on log(size) coefficient |
| $\gamma_{{size1}_{s}}$ | Parameter | Random species effect on log(size) coefficient |
| $\beta_{{size2}_{s}}$ | Transformed parameter | Size coefficient |
| $\beta_{size2}$ | Parameter | Average size coefficient |
| $\boldsymbol{\delta}_{\mathbf{size2}}$ (3) | Parameter | Trait effects on size coefficient |
| $\gamma_{{size2}_{s}}$ | Parameter | Random species effect on size coefficient |
| ${crowding}_{is}$ | Transformed parameter | Crowding term |
| $\mathrm{BA}_{i}$ | Data | Neighbor basal area (cm^2^) |
| ${\beta_{\mathrm{crowd}}}_{s}$ | Transformed parameter | Crowding response |
| $\beta_{\mathrm{crowd}}$ | Parameter | Average crowding response |
| $\gamma_{\mathrm{crowd}_{s}}$ | Parameter | Random species effect on crowding response |
| ${climate}_{sp}$ | Transformed parameter | Climate term |
| $\mathrm{MAT}_{p}$ | Data | Mean annual temperature (°C) |
| $\beta_{\mathrm{clim}_{s}}$ | Transformed parameter | Temperature response |
| $\beta_{\mathrm{clim}}$ | Parameter | Average temperature response |
| $\gamma_{\mathrm{clim}_{s}}$ | Parameter | Random species effect on temperature response |
| $trait_{sp}$ | Transformed parameter | Trait term (performance landscape) |
| $\mathbf{trait}_{s}$ (3) | Data | Trait vector (wood density, SLA, maximum height) |
| $\boldsymbol{\beta}_{\mathbf{dir}_{p}}$ (3) | Transformed parameter | Directional (linear) performance gradients |
| $\beta_{\mathrm{dir}_{t}}$ | Parameter | Average directional performance gradient |
| $\delta_{\mathrm{dir}_{t}}$ | Parameter | Temperature effect on directional performance gradient |
| $\boldsymbol{\beta}_{\mathbf{nonlin}_{p}}$ (3x3) | Transformed parameter | Nonlinear performance gradients |
| $\beta_{\mathrm{nonlin}_{tu}}$ | Parameter | Average nonlinear performance gradient |
| $\delta_{\mathrm{nonlin}_{tu}}$ | Parameter | Temperature effect on nonlinear performance gradient |
| $\tau_{\mathrm{plot}}$ | Parameter | Plot random effects standard deviation |
| $\boldsymbol{\Sigma}$ (5x5) | Transformed parameter | Species random effects covariance matrix |
| $\boldsymbol{\tau}_{\mathbf{species}}$ (5) | Parameter | Species random effects standard deviations |
| ***L*** (5x5) | Parameter | Species random effects correlation matrix (Cholesky factor) |

**Table S4.** Descriptions of terms and variables in the recruitment model. Terms are categorized as data (observed quantities), parameters (unobserved quantities that are directly sampled), and transformed parameters (unobserved quantities that are functions of parameters and data). Dimensions of vector and matrix terms are given in parentheses. Subscripts reference individual *i*, species *s*, plot *p*, trait *t* and *u*, observation (tree-year) *j*, tree *q*.

| **Term** | **Type** | **Description** |
| --- | --- | --- |
| $\mathrm{recruitment}_{sp}$ | Data (response variable) | Observed plot-level recruitment (number of recruits) |
| ${recr}_{sp}$ | Transformed parameter | Expected plot-level recruitment rate (recruits year^-1^) |
| ${RI}_{p}$ | Data | Remeasurement interval (years) |
| $\phi$ | Parameter | Overdispersion of negative binomial distribution |
| $\mathrm{repro}_{jqsp}$ | Data | Observed reproductive status (binary) |
| $pr_{jqsp}$ | Transformed parameter | Expected probability of being reproductive |
| $\gamma_{{site}_{p}}$ | Parameter | Random site effect on reproductive status |
| $\gamma_{\mathrm{tree}_{q}}$ | Parameter | Random tree effect on reproductive status |
| $R_{isp}$ | Transformed parameter | Expected per-capita recruitment rate (recruits year^-1^) |
| ${PR}_{sp}$ | Transformed parameter | Potential per-capita recruitment rate |
| $\alpha_{\mathrm{PR}}$ | Parameter | Average potential recruitment rate |
| $\gamma_{\mathrm{PR}_{s}}$ | Parameter | Random species effect on potential recruitment rate |
| $\gamma_{\mathrm{PR}_{p}}$ | Parameter | Random plot effect on potential recruitment rate |
| ${size}_{is}$ | Transformed parameter | Size term |
| $\mathrm{dia}_{i}$ | Data | Diameter at breast height (cm/30) |
| $D_{s}$ | Transformed parameter | Reproductive size threshold (log(cm/30) |
| $D$ | Parameter | Average reproductive size threshold |
| $\boldsymbol{\delta}_{\mathbf{D}}$ (3) | Parameter | Trait effects on reproductive size threshold |
| $\gamma_{D_{s}}$ | Parameter | Random species effect on reproductive size threshold |
| ${\beta_{\mathrm{repro}}}_{s}$ | Transformed parameter | Slope of size effect on reproductive status |
| $\beta_{\mathrm{repro}}$ | Parameter | Average slope of size effect on reproductive status |
| $\boldsymbol{\delta}_{\mathbf{repro}}$ (3) | Parameter | Trait effects on slope of size-reproductive status effect |
| $\gamma_{\mathrm{repro}_{s}}$ | Parameter | Random species effect on slope of size-reproductive status effect |
| $\beta_{\mathrm{recr}_{s}}$ | Transformed parameter | Size effect on per-capita recruitment |
| $\beta_{\mathrm{recr}}$ | Parameter | Average size effect on per-capita recruitment |
| $\boldsymbol{\delta}_{\mathbf{recr}}$ (3) | Parameter | Trait effects on size-recruitment effect |
| $\gamma_{\mathrm{recr}_{s}}$ | Parameter | Random species effect on size-recruitment effect |
| ${crowding}_{is}$ | Transformed parameter | Crowding term |
| $\mathrm{BA}_{\mathrm{can}_{i}}$ | Data | Canopy neighbor basal area (cm^2^) |
| ${\beta_{\mathrm{can}}}_{s}$ | Transformed parameter | Response to crowding by canopy trees |
| $\beta_{\mathrm{can}}$ | Parameter | Average response to crowding by canopy trees |
| $\gamma_{\mathrm{can}_{s}}$ | Parameter | Random species effect on canopy crowding response |
| $\mathrm{BA}_{\mathrm{sap}_{i}}$ | Data | Sapling neighbor basal area (cm^2^) |
| ${\beta_{\mathrm{sap}}}_{s}$ | Transformed parameter | Response to crowding by saplings |
| $\beta_{\mathrm{sap}}$ | Parameter | Average response to crowding by saplings |
| $\gamma_{\mathrm{sap}_{s}}$ | Parameter | Random species effect on sapling crowding response |

**Table S4** continued.

| **Term** | **Type** | **Description** |
| --- | --- | --- |
| ${climate}_{sp}$ | Transformed parameter | Climate term |
| $\mathrm{MAT}_{p}$ | Data | Mean annual temperature (°C) |
| $\beta_{\mathrm{clim}_{s}}$ | Transformed parameter | Temperature response |
| $\beta_{\mathrm{clim}}$ | Parameter | Average temperature response |
| $\gamma_{\mathrm{clim}_{s}}$ | Parameter | Random species effect on temperature response |
| $trait_{sp}$ | Transformed parameter | Trait term (performance landscape) |
| $\mathbf{trait}_{s}$ (3) | Data | Trait vector (wood density, SLA, maximum height) |
| $\boldsymbol{\beta}_{\mathbf{dir}_{p}}$ (3) | Transformed parameter | Directional (linear) performance gradients |
| $\beta_{\mathrm{dir}_{t}}$ | Parameter | Average directional performance gradient |
| $\delta_{\mathrm{dir}_{t}}$ | Parameter | Temperature effect on directional performance gradient |
| $\boldsymbol{\beta}_{\mathbf{nonlin}_{p}}$ (3x3) | Transformed parameter | Nonlinear performance gradients |
| $\beta_{\mathrm{nonlin}_{tu}}$ | Parameter | Average nonlinear performance gradient |
| $\delta_{\mathrm{nonlin}_{tu}}$ | Parameter | Temperature effect on nonlinear performance gradient |
| $\tau_{\mathrm{plot}}$ | Parameter | Plot random effects standard deviation |
| $\boldsymbol{\Sigma}$ (7x7) | Transformed parameter | Species random effects covariance matrix |
| $\boldsymbol{\tau}_{\mathbf{species}}$ (7) | Parameter | Species random effects standard deviations |
| ***L*** (7x7) | Parameter | Species random effects correlation matrix (Cholesky factor) |
| $\tau_{\mathrm{site}}$ | Parameter | Standard deviation of site random effects |
| $\tau_{\mathrm{tree}}$ | Parameter | Standard deviation of tree random effects |

**Table S5.** Priors for demographic rate models.

| Model | Parameter | Prior |
| --- | --- | --- |
| Sapling survival | *α*_PS_ | Normal(0, 1) |
|  | *β*_size1_ | Normal(0, 1) |
|  | ***δ*_size1_** | Normal(0, 1) |
|  | *β*_crowd_ | Normal(0, 1) |
|  | *β*_clim_ | Normal(0, 1) |
|  | ***β*_dir_** | Normal(0, 1) |
|  | ***β*_nonlin_** | Normal(0, 1) |
|  | ***δ*_dir_** | Normal(0, 1) |
|  | ***δ*_nonlin_** | Normal(0, 1) |
|  | *τ*_plot_ | Normal(0, 0.25) |
|  | ***τ*_species_** | Normal(0, 0.25) |
|  | ***L*** | LkjCorr(2) |
| Canopy tree survival | *α*_PS_ | Normal(0, 1) |
|  | *β*_size1,_ *β*_size2_ | Normal(0, 1) |
|  | ***δ*_size1,_ *δ*_size2_** | Normal(0, 1) |
|  | *β*_crowd_ | Normal(0, 1) |
|  | *β*_clim_ | Normal(0, 1) |
|  | ***β*_dir_** | Normal(0, 1) |
|  | ***β*_nonlin_** | Normal(0, 1) |
|  | ***δ*_dir_** | Normal(0, 1) |
|  | ***δ*_nonlin_** | Normal(0, 1) |
|  | *τ*_plot_ | Normal(0, 0.25) |
|  | *τ*_species_ | Normal(0, 0.25) |
|  | ***L*** | LkjCorr(2) |
| Growth | *α*_PG_ | Normal(0, 1) |
|  | *β*_size1_, *β*_size2_ | Normal(0, 1) |
|  | ***δ*_size1_, *δ*_size2_** | Normal(0, 1) |
|  | *β*_crowd_ | Normal(0, 1) |
|  | *β*_clim_ | Normal(0, 1) |
|  | ***β*_dir_** | Normal(0, 1) |
|  | ***β*_nonlin_** | Normal(0, 1) |
|  | ***δ*_dir_** | Normal(0, 1) |
|  | ***δ*_nonlin_** | Normal(0, 1) |
|  | *τ*_plot_ | Normal(0, 0.25) |
|  | ***τ*_species_** | Normal(0, 0.25) |
|  | ***L*** | LkjCorr(2) |
|  | *β* | LogNormal(0.75, 0.75) |
|  | *z* | Normal(0, 1) |

**Table S5** continued.

| Model | Parameter | Prior |
| --- | --- | --- |
| Recruitment | *α*_PR_ | Normal(-1, 1) |
|  | *D* | Normal(0, 1) |
|  | ***δ*_D_** | Normal(0, 1) |
|  | *β*_repro_ | Normal(0, 1) |
|  | ***δ*_repro_** | Normal(0, 1) |
|  | *β*_recr_ | Normal(-0.5, 0.5) |
|  | ***δ*_recr_** | Normal(0, 0.2) |
|  | *β*_can_ | Normal(0, 1) |
|  | *β*_sap_ | Normal(0, 1) |
|  | *β*_clim_ | Normal(0, 1) |
|  | ***β*_dir_** | Normal(0, 1) |
|  | ***β*_nonlin_** | Normal(0, 1) |
|  | ***δ*_dir_** | Normal(0, 1) |
|  | ***δ*_nonlin_** | Normal(0, 1) |
|  | *τ*_plot_ | Normal(0, 0.2) |
|  | ***τ*_species_** | Normal(0, 0.2) |
|  | *τ*_site_ | Normal(0, 0.2) |
|  | *τ*_tree_ | Normal(0, 0.2) |
|  | ***L*** | LkjCorr(2) |
|  | *Φ*^-0.5^ | Normal(0, 1) |

**Table S6.** Sapling survival parameter estimates and convergence statistic.

| Parameter | Mean | SD | 5% | 50% | 95% | $\hat{R}$ |
| --- | --- | --- | --- | --- | --- | --- |
| $\alpha_{\mathrm{PS}}$ | 3.26 | 0.17 | 2.97 | 3.26 | 3.53 | 1.00 |
| $\beta_{\mathrm{clim}}$ | -0.01 | 0.09 | -0.15 | -0.01 | 0.14 | 1.00 |
| $\beta_{size1}$ | 0.47 | 0.03 | 0.42 | 0.47 | 0.52 | 1.00 |
| $\beta_{\mathrm{crowd}}$ | -0.07 | 0.02 | -0.11 | -0.07 | -0.03 | 1.00 |
| $\beta_{\mathrm{dir}_{WD}}$ | 0.14 | 0.11 | -0.04 | 0.14 | 0.32 | 1.00 |
| $\beta_{\mathrm{dir}_{SLA}}$ | 0.11 | 0.11 | -0.07 | 0.11 | 0.29 | 1.00 |
| $\beta_{\mathrm{dir}_{Ht}}$ | 0.19 | 0.11 | 0.00 | 0.19 | 0.38 | 1.00 |
| $\beta_{\mathrm{nonlin}_{WD}}$ | 0.00 | 0.09 | -0.15 | 0.00 | 0.14 | 1.00 |
| $\beta_{\mathrm{nonlin}_{SLA}}$ | 0.13 | 0.09 | -0.01 | 0.13 | 0.29 | 1.00 |
| $\beta_{\mathrm{nonlin}_{Ht}}$ | -0.10 | 0.08 | -0.22 | -0.10 | 0.03 | 1.00 |
| $\beta_{\mathrm{nonlin}_{WD-SLA}}$ | 0.24 | 0.10 | 0.07 | 0.24 | 0.41 | 1.00 |
| $\beta_{\mathrm{nonlin}_{WD-Ht}}$ | -0.03 | 0.09 | -0.18 | -0.03 | 0.12 | 1.00 |
| $\beta_{\mathrm{nonlin}_{SLA-Ht}}$ | 0.11 | 0.10 | -0.06 | 0.12 | 0.28 | 1.00 |
| $\delta_{{size1}_{WD}}$ | -0.01 | 0.03 | -0.07 | -0.01 | 0.04 | 1.00 |
| $\delta_{{size1}_{SLA}}$ | -0.04 | 0.03 | -0.10 | -0.04 | 0.01 | 1.00 |
| $\delta_{{size1}_{Ht}}$ | 0.00 | 0.03 | -0.05 | 0.00 | 0.05 | 1.00 |
| $\delta_{\mathrm{dir}_{WD}}$ | 0.03 | 0.06 | -0.07 | 0.03 | 0.12 | 1.00 |
| $\delta_{\mathrm{dir}_{SLA}}$ | -0.02 | 0.06 | -0.11 | -0.02 | 0.08 | 1.00 |
| $\delta_{\mathrm{dir}_{Ht}}$ | 0.00 | 0.06 | -0.10 | 0.00 | 0.10 | 1.00 |
| $\delta_{\mathrm{nonlin}_{WD}}$ | 0.02 | 0.05 | -0.06 | 0.02 | 0.11 | 1.00 |
| $\delta_{\mathrm{nonlin}_{SLA}}$ | -0.04 | 0.05 | -0.12 | -0.04 | 0.04 | 1.00 |
| $\delta_{\mathrm{nonlin}_{Ht}}$ | -0.02 | 0.04 | -0.09 | -0.02 | 0.05 | 1.00 |
| $\delta_{\mathrm{nonlin}_{WD-SLA}}$ | 0.06 | 0.06 | -0.04 | 0.06 | 0.16 | 1.00 |
| $\delta_{\mathrm{nonlin}_{WD-Ht}}$ | -0.03 | 0.04 | -0.11 | -0.03 | 0.04 | 1.00 |
| $\delta_{\mathrm{nonlin}_{SLA-Ht}}$ | 0.04 | 0.05 | -0.04 | 0.04 | 0.13 | 1.00 |
| $\tau_{\mathrm{plot}}$ | 0.73 | 0.02 | 0.70 | 0.73 | 0.77 | 1.01 |
| $\tau_{\mathrm{spe}cies[\gamma_{\mathrm{PS}}]}$ | 0.74 | 0.07 | 0.62 | 0.73 | 0.86 | 1.00 |
| $\tau_{\mathrm{spe}cies[\gamma_{\mathrm{clim}}]}$ | 0.23 | 0.06 | 0.14 | 0.23 | 0.33 | 1.00 |
| $\tau_{\mathrm{spe}cies[\gamma_{size1}]}$ | 0.19 | 0.03 | 0.14 | 0.19 | 0.24 | 1.00 |
| $\tau_{\mathrm{spe}cies[\gamma_{\mathrm{crowd}}]}$ | 0.13 | 0.03 | 0.09 | 0.13 | 0.17 | 1.00 |

**Table S7.** Canopy tree survival model parameter estimates and convergence statistic.

| Parameter | Mean | SD | 5% | 50% | 95% | $\hat{R}$ |
| --- | --- | --- | --- | --- | --- | --- |
| $\alpha_{\mathrm{PS}}$ | 4.07 | 0.16 | 3.80 | 4.07 | 4.33 | 1.00 |
| $\beta_{\mathrm{clim}}$ | 0.12 | 0.13 | -0.08 | 0.11 | 0.33 | 1.00 |
| $\beta_{size2}$ | -0.48 | 0.07 | -0.60 | -0.48 | -0.36 | 1.00 |
| $\beta_{size1}$ | 0.65 | 0.07 | 0.53 | 0.65 | 0.77 | 1.00 |
| $\beta_{\mathrm{crowd}}$ | -0.19 | 0.02 | -0.22 | -0.19 | -0.15 | 1.00 |
| $\beta_{\mathrm{dir}_{WD}}$ | 0.33 | 0.10 | 0.17 | 0.33 | 0.50 | 1.00 |
| $\beta_{\mathrm{dir}_{SLA}}$ | 0.09 | 0.10 | -0.08 | 0.09 | 0.25 | 1.00 |
| $\beta_{\mathrm{dir}_{Ht}}$ | 0.45 | 0.11 | 0.27 | 0.46 | 0.62 | 1.00 |
| $\beta_{\mathrm{nonlin}_{WD}}$ | 0.11 | 0.09 | -0.04 | 0.11 | 0.27 | 1.00 |
| $\beta_{\mathrm{nonlin}_{SLA}}$ | 0.10 | 0.09 | -0.04 | 0.10 | 0.25 | 1.00 |
| $\beta_{\mathrm{nonlin}_{Ht}}$ | -0.05 | 0.08 | -0.18 | -0.06 | 0.08 | 1.00 |
| $\beta_{\mathrm{nonlin}_{WD-SLA}}$ | 0.02 | 0.10 | -0.15 | 0.02 | 0.19 | 1.00 |
| $\beta_{\mathrm{nonlin}_{WD-Ht}}$ | 0.03 | 0.09 | -0.13 | 0.03 | 0.18 | 1.00 |
| $\beta_{\mathrm{nonlin}_{SLA-Ht}}$ | 0.14 | 0.10 | -0.01 | 0.14 | 0.30 | 1.00 |
| $\delta_{{size2}_{WD}}$ | -0.10 | 0.06 | -0.20 | -0.10 | 0.01 | 1.00 |
| $\delta_{{size2}_{SLA}}$ | 0.17 | 0.07 | 0.06 | 0.17 | 0.28 | 1.00 |
| $\delta_{{size2}_{Ht}}$ | -0.03 | 0.08 | -0.16 | -0.02 | 0.11 | 1.00 |
| $\delta_{{size1}_{WD}}$ | 0.11 | 0.06 | 0.01 | 0.11 | 0.22 | 1.00 |
| $\delta_{{size1}_{SLA}}$ | -0.19 | 0.07 | -0.30 | -0.19 | -0.08 | 1.00 |
| $\delta_{{size1}_{Ht}}$ | 0.15 | 0.08 | 0.02 | 0.15 | 0.28 | 1.00 |
| $\delta_{\mathrm{dir}_{WD}}$ | 0.06 | 0.07 | -0.06 | 0.06 | 0.18 | 1.00 |
| $\delta_{\mathrm{dir}_{SLA}}$ | 0.06 | 0.07 | -0.06 | 0.06 | 0.17 | 1.00 |
| $\delta_{\mathrm{dir}_{Ht}}$ | -0.11 | 0.08 | -0.25 | -0.11 | 0.02 | 1.00 |
| $\delta_{\mathrm{nonlin}_{WD}}$ | 0.03 | 0.07 | -0.08 | 0.03 | 0.14 | 1.00 |
| $\delta_{\mathrm{nonlin}_{SLA}}$ | -0.07 | 0.07 | -0.18 | -0.07 | 0.04 | 1.00 |
| $\delta_{\mathrm{nonlin}_{Ht}}$ | -0.08 | 0.06 | -0.18 | -0.08 | 0.02 | 1.00 |
| $\delta_{\mathrm{nonlin}_{WD-SLA}}$ | -0.06 | 0.08 | -0.18 | -0.06 | 0.06 | 1.00 |
| $\delta_{\mathrm{nonlin}_{WD-Ht}}$ | -0.06 | 0.06 | -0.17 | -0.06 | 0.04 | 1.00 |
| $\delta_{\mathrm{nonlin}_{SLA-Ht}}$ | 0.04 | 0.07 | -0.07 | 0.04 | 0.16 | 1.00 |
| $\tau_{\mathrm{plot}}$ | 0.68 | 0.01 | 0.66 | 0.68 | 0.70 | 1.00 |
| $\tau_{species[\gamma_{\mathrm{PS}}]}$ | 0.80 | 0.07 | 0.68 | 0.79 | 0.92 | 1.00 |
| $\tau_{species[\gamma_{\mathrm{clim}}]}$ | 0.51 | 0.06 | 0.42 | 0.51 | 0.61 | 1.00 |
| $\tau_{species[\gamma_{size2}]}$ | 0.23 | 0.07 | 0.12 | 0.23 | 0.35 | 1.00 |
| $\tau_{species[\gamma_{size1}]}$ | 0.33 | 0.06 | 0.24 | 0.32 | 0.43 | 1.01 |
| $\tau_{species[\gamma_{\mathrm{crowd}}]}$ | 0.13 | 0.02 | 0.10 | 0.13 | 0.16 | 1.00 |

**Table S8.** Growth model parameter estimates and convergence statistic.

| Parameter | Mean | SD | 5% | 50% | 95% | $\hat{R}$ |
| --- | --- | --- | --- | --- | --- | --- |
| $\alpha_{\mathrm{PG}}$ | 0.89 | 0.05 | 0.82 | 0.89 | 0.97 | 1.03 |
| $\beta_{\mathrm{clim}}$ | 0.00 | 0.03 | -0.05 | 0.00 | 0.06 | 1.01 |
| $\beta_{size2}$ | 0.39 | 0.02 | 0.36 | 0.39 | 0.42 | 1.00 |
| $\beta_{size1}$ | -0.10 | 0.02 | -0.13 | -0.10 | -0.07 | 1.00 |
| $\beta_{\mathrm{crowd}}$ | -0.13 | 0.01 | -0.14 | -0.13 | -0.11 | 1.00 |
| $\beta_{\mathrm{dir}_{WD}}$ | -0.04 | 0.03 | -0.09 | -0.04 | 0.01 | 1.02 |
| $\beta_{\mathrm{dir}_{SLA}}$ | 0.03 | 0.03 | -0.02 | 0.03 | 0.08 | 1.01 |
| $\beta_{\mathrm{dir}_{Ht}}$ | 0.12 | 0.03 | 0.06 | 0.12 | 0.17 | 1.01 |
| $\beta_{\mathrm{nonlin}_{WD}}$ | -0.01 | 0.02 | -0.05 | -0.01 | 0.03 | 1.01 |
| $\beta_{\mathrm{nonlin}_{SLA}}$ | -0.04 | 0.03 | -0.08 | -0.04 | 0.00 | 1.02 |
| $\beta_{\mathrm{nonlin}_{Ht}}$ | -0.01 | 0.02 | -0.04 | -0.01 | 0.03 | 1.00 |
| $\beta_{\mathrm{nonlin}_{WD-SLA}}$ | -0.02 | 0.03 | -0.07 | -0.02 | 0.03 | 1.01 |
| $\beta_{\mathrm{nonlin}_{WD-Ht}}$ | 0.00 | 0.03 | -0.04 | 0.00 | 0.04 | 1.01 |
| $\beta_{\mathrm{nonlin}_{SLA-Ht}}$ | -0.04 | 0.03 | -0.08 | -0.03 | 0.01 | 1.00 |
| $\delta_{{size2}_{WD}}$ | 0.00 | 0.02 | -0.03 | 0.00 | 0.03 | 1.00 |
| $\delta_{{size2}_{SLA}}$ | -0.02 | 0.02 | -0.06 | -0.02 | 0.01 | 1.01 |
| $\delta_{{size2}_{Ht}}$ | 0.03 | 0.02 | 0.00 | 0.03 | 0.07 | 1.00 |
| $\delta_{{size1}_{WD}}$ | 0.03 | 0.02 | 0.00 | 0.03 | 0.05 | 1.01 |
| $\delta_{{size1}_{SLA}}$ | 0.01 | 0.02 | -0.02 | 0.01 | 0.04 | 1.00 |
| $\delta_{{size1}_{Ht}}$ | 0.01 | 0.02 | -0.02 | 0.01 | 0.04 | 1.00 |
| $\delta_{\mathrm{dir}_{WD}}$ | 0.02 | 0.02 | -0.02 | 0.02 | 0.05 | 1.01 |
| $\delta_{\mathrm{dir}_{SLA}}$ | -0.04 | 0.02 | -0.07 | -0.04 | 0.00 | 1.01 |
| $\delta_{\mathrm{dir}_{Ht}}$ | -0.03 | 0.02 | -0.06 | -0.03 | 0.01 | 1.00 |
| $\delta_{\mathrm{nonlin}_{WD}}$ | -0.01 | 0.02 | -0.04 | -0.01 | 0.01 | 1.02 |
| $\delta_{\mathrm{nonlin}_{SLA}}$ | 0.01 | 0.02 | -0.02 | 0.01 | 0.04 | 1.01 |
| $\delta_{\mathrm{nonlin}_{Ht}}$ | 0.02 | 0.02 | -0.01 | 0.02 | 0.04 | 1.00 |
| $\delta_{\mathrm{nonlin}_{WD-SLA}}$ | -0.02 | 0.02 | -0.05 | -0.02 | 0.01 | 1.01 |
| $\delta_{\mathrm{nonlin}_{WD-Ht}}$ | 0.01 | 0.02 | -0.02 | 0.01 | 0.04 | 1.01 |
| $\delta_{\mathrm{nonlin}_{SLA-Ht}}$ | 0.01 | 0.02 | -0.02 | 0.01 | 0.04 | 1.00 |
| $\tau_{\mathrm{plot}}$ | 0.25 | 0.00 | 0.24 | 0.25 | 0.25 | 1.00 |
| $\tau_{species[\gamma_{\mathrm{PS}}]}$ | 0.24 | 0.02 | 0.21 | 0.24 | 0.28 | 1.01 |
| $\tau_{species[\gamma_{\mathrm{clim}}]}$ | 0.15 | 0.02 | 0.12 | 0.14 | 0.17 | 1.01 |
| $\tau_{species[\gamma_{size2}]}$ | 0.17 | 0.02 | 0.14 | 0.17 | 0.20 | 1.00 |
| $\tau_{species[\gamma_{size1}]}$ | 0.13 | 0.01 | 0.11 | 0.13 | 0.16 | 1.01 |
| $\tau_{species[\gamma_{\mathrm{crowd}}]}$ | 0.07 | 0.01 | 0.05 | 0.06 | 0.08 | 1.00 |
| $\beta$ | 1.00 | 1.00 | 1.01 | 1.00 | 0.00 | 1.00 |
| *z* | -1.63 | -1.62 | -1.61 | -1.62 | 0.01 | 1.00 |

**Table S9.** Recruitment model parameter estimates and convergence statistic.

| Parameter | Mean | SD | 5% | 50% | 95% | $\hat{R}$ |
| --- | --- | --- | --- | --- | --- | --- |
| $\alpha_{\mathrm{PR}}$ | -4.53 | 0.23 | -4.92 | -4.53 | -4.16 | 1.00 |
| $D$ | -1.48 | 0.12 | -1.68 | -1.48 | -1.29 | 1.02 |
| $\beta_{\mathrm{repro}}$ | 1.59 | 0.06 | 1.49 | 1.59 | 1.69 | 1.00 |
| $\beta_{\mathrm{recr}}$ | -1.92 | 0.28 | -2.40 | -1.91 | -1.48 | 1.01 |
| $\delta_{D_{WD}}$ | 0.23 | 0.08 | 0.10 | 0.23 | 0.36 | 1.01 |
| $\delta_{D_{SLA}}$ | -0.05 | 0.07 | -0.17 | -0.05 | 0.07 | 1.00 |
| $\delta_{D_{Ht}}$ | 0.43 | 0.08 | 0.30 | 0.43 | 0.56 | 1.00 |
| $\delta_{\mathrm{repro}_{WD}}$ | 0.06 | 0.05 | -0.03 | 0.06 | 0.15 | 1.00 |
| $\delta_{\mathrm{repro}_{SLA}}$ | -0.02 | 0.05 | -0.11 | -0.02 | 0.06 | 1.00 |
| $\delta_{\mathrm{repro}_{Ht}}$ | 0.18 | 0.05 | 0.10 | 0.18 | 0.26 | 1.00 |
| $\delta_{\mathrm{recr}_{WD}}$ | 0.16 | 0.15 | -0.09 | 0.16 | 0.41 | 1.00 |
| $\delta_{\mathrm{recr}_{SLA}}$ | 0.15 | 0.16 | -0.11 | 0.16 | 0.41 | 1.00 |
| $\delta_{\mathrm{recr}_{Ht}}$ | -0.04 | 0.15 | -0.28 | -0.04 | 0.21 | 1.00 |
| $\beta_{\mathrm{clim}}$ | 0.07 | 0.18 | -0.23 | 0.08 | 0.37 | 1.00 |
| $\beta_{\mathrm{can}}$ | -0.49 | 0.10 | -0.66 | -0.48 | -0.32 | 1.00 |
| $\beta_{\mathrm{sap}}$ | -0.44 | 0.09 | -0.59 | -0.43 | -0.30 | 1.00 |
| $\beta_{\mathrm{dir}_{WD}}$ | -0.38 | 0.13 | -0.59 | -0.38 | -0.17 | 1.00 |
| $\beta_{\mathrm{dir}_{SLA}}$ | 0.22 | 0.13 | 0.01 | 0.21 | 0.43 | 1.00 |
| $\beta_{\mathrm{dir}_{Ht}}$ | 0.00 | 0.13 | -0.22 | -0.01 | 0.21 | 1.00 |
| $\beta_{\mathrm{nonlin}_{WD}}$ | -0.11 | 0.11 | -0.29 | -0.11 | 0.07 | 1.00 |
| $\beta_{\mathrm{nonlin}_{SLA}}$ | -0.01 | 0.10 | -0.17 | -0.01 | 0.16 | 1.00 |
| $\beta_{\mathrm{nonlin}_{Ht}}$ | 0.08 | 0.09 | -0.07 | 0.08 | 0.23 | 1.00 |
| $\beta_{\mathrm{nonlin}_{WD-SLA}}$ | 0.42 | 0.13 | 0.21 | 0.42 | 0.64 | 1.00 |
| $\beta_{\mathrm{nonlin}_{WD-Ht}}$ | 0.10 | 0.10 | -0.07 | 0.10 | 0.27 | 1.00 |
| $\beta_{\mathrm{nonlin}_{SLA-Ht}}$ | -0.03 | 0.11 | -0.21 | -0.03 | 0.15 | 1.00 |
| $\delta_{\mathrm{dir}_{WD}}$ | -0.13 | 0.10 | -0.30 | -0.13 | 0.04 | 1.00 |
| $\delta_{\mathrm{dir}_{SLA}}$ | 0.07 | 0.10 | -0.10 | 0.07 | 0.23 | 1.00 |
| $\delta_{\mathrm{dir}_{Ht}}$ | -0.14 | 0.11 | -0.33 | -0.14 | 0.04 | 1.00 |
| $\delta_{\mathrm{nonlin}_{WD}}$ | 0.13 | 0.10 | -0.02 | 0.13 | 0.30 | 1.00 |
| $\delta_{\mathrm{nonlin}_{SLA}}$ | -0.13 | 0.08 | -0.27 | -0.13 | 0.01 | 1.00 |
| $\delta_{\mathrm{nonlin}_{Ht}}$ | 0.06 | 0.08 | -0.07 | 0.06 | 0.20 | 1.00 |
| $\delta_{\mathrm{nonlin}_{WD-SLA}}$ | 0.09 | 0.11 | -0.09 | 0.09 | 0.26 | 1.00 |
| $\delta_{\mathrm{nonlin}_{WD-Ht}}$ | 0.07 | 0.08 | -0.07 | 0.07 | 0.21 | 1.00 |
| $\delta_{\mathrm{nonlin}_{SLA-Ht}}$ | 0.07 | 0.09 | -0.08 | 0.06 | 0.21 | 1.00 |
| $\tau_{\mathrm{plot}}$ | 0.84 | 0.04 | 0.77 | 0.84 | 0.91 | 1.00 |
| $\tau_{species[\gamma_{\mathrm{PR}}]}$ | 0.48 | 0.09 | 0.34 | 0.48 | 0.63 | 1.00 |
| $\tau_{species[\gamma_{D}]}$ | 1.33 | 0.12 | 1.13 | 1.32 | 1.53 | 1.00 |
| $\tau_{species[\gamma_{\mathrm{repro}}]}$ | 0.29 | 0.24 | 0.02 | 0.23 | 0.78 | 1.01 |
| $\tau_{species[\gamma_{\mathrm{recr}}]}$ | 0.42 | 0.17 | 0.09 | 0.44 | 0.69 | 1.00 |

**Table S9** continued.

| Parameter | Mean | SD | 5% | 50% | 95% | $\hat{R}$ |
| --- | --- | --- | --- | --- | --- | --- |
| $\tau_{species[\gamma_{\mathrm{clim}}]}$ | 0.35 | 0.08 | 0.21 | 0.34 | 0.49 | 1.00 |
| $\tau_{species[\gamma_{\mathrm{can}}]}$ | 0.20 | 0.07 | 0.08 | 0.20 | 0.32 | 1.00 |
| $\tau_{species[\gamma_{\mathrm{sap}}]}$ | 0.20 | 0.07 | 0.10 | 0.19 | 0.31 | 1.01 |
| $\tau_{\mathrm{site}}$ | 1.35 | 0.12 | 1.16 | 1.35 | 1.55 | 1.00 |
| $\tau_{\mathrm{tree}}$ | 1.98 | 0.08 | 1.84 | 1.98 | 2.12 | 1.01 |
| $\phi$ | 0.54 | 0.45 | 0.60 | 0.54 | 0.49 | 1.00 |

**Table S10.** List of tree species included in demographic rate models. WD, wood density (g cm^-3^); SLA, specific leaf area (mm^2^ mg^-1^); HMax, maximum height (m).

|  |  |  |  |  | Included in model? | | | | |
| --- | --- | --- | --- | --- | --- | --- | --- | --- | --- |
| Species | Abbrev. | WD | SLA | HMax | Sapling survival | Canopy survival | Growth | Recruit- ment | Repro. status |
| *Abies balsamea* | *Abi.bal* | 0.42 | 5.6 | 23.2 | Y | Y | Y | Y | Y |
| *Acer negundo* | *Ace.neg* | 0.53 | 24.1 | 19.5 | Y | Y | Y | Y | Y |
| *Acer pensylvanicum* | *Ace.pen* | 0.44 | 28.1 | 11.7 | Y | Y | Y | Y | Y |
| *Acer platanoides* | *Ace.pla* | 0.47 | 20.8 | 29.5 | N | Y | Y | N | N |
| *Acer rubrum* | *Ace.rub* | 0.52 | 19.9 | 29.1 | Y | Y | Y | Y | Y |
| *Acer saccharinum* | *Ace.san* | 0.48 | 21.2 | 30.3 | N | Y | Y | N | N |
| *Acer saccharum* | *Ace.sac* | 0.62 | 22.1 | 30.2 | Y | Y | Y | Y | Y |
| *Aesculus flava* | *Aes.fla* | 0.35 | 26.6 | 29.8 | Y | Y | Y | Y | Y |
| *Ailanthus altissima* | *Ail.alt* | 0.53 | 31.8 | 22.6 | Y | Y | Y | Y | Y |
| *Betula alleghaniensis* | *Bet.all* | 0.61 | 25.5 | 29.1 | Y | Y | Y | Y | Y |
| *Betula lenta* | *Bet.len* | 0.66 | 25.0 | 23.9 | Y | Y | Y | Y | Y |
| *Betula nigra* | *Bet.nig* | 0.50 | 15.8 | 24.0 | Y | Y | Y | Y | N |
| *Betula papyrifera* | *Bet.pap* | 0.54 | 18.4 | 25.4 | Y | Y | Y | Y | N |
| *Betula populifolia* | *Bet.pop* | 0.50 | 22.2 | 10.8 | Y | Y | Y | Y | N |
| *Carpinus caroliniana* | *Car.car* | 0.65 | 26.8 | 12.9 | Y | Y | Y | Y | Y |
| *Carya alba* | *Car.alb* | 0.69 | 20.8 | 23.3 | Y | Y | Y | Y | Y |
| *Carya cordiformis* | *Car.cor* | 0.59 | 28.3 | 39.0 | Y | Y | Y | Y | N |
| *Carya glabra* | *Car.gla* | 0.71 | 18.3 | 28.8 | Y | Y | Y | Y | Y |
| *Carya illinoinensis* | *Car.ill* | 0.65 | 22.4 | 38.9 | N | Y | Y | N | N |
| *Carya laciniosa* | *Car.lac* | 0.64 | 22.4 | 31.7 | N | Y | Y | Y | N |
| *Carya ovata* | *Car.ova* | 0.69 | 16.9 | 35.4 | Y | Y | Y | Y | Y |
| *Castanea dentata* | *Cas.den* | 0.45 | 19.4 | 20.0 | Y | Y | Y | Y | N |
| *Celtis laevigata* | *Cel.lae* | 0.47 | 19.9 | 24.2 | Y | Y | Y | Y | Y |
| *Celtis occidentalis* | *Cel.occ* | 0.49 | 22.9 | 21.2 | Y | Y | Y | Y | N |
| *Cercis canadensis* | *Cer.can* | 0.65 | 25.2 | 10.9 | Y | Y | Y | Y | Y |
| *Cornus florida* | *Cor.flo* | 0.68 | 19.5 | 12.2 | Y | Y | Y | Y | Y |
| *Diospyros virginiana* | *Dio.vir* | 0.65 | 23.6 | 18.6 | Y | Y | Y | Y | Y |
| *Fagus grandifolia* | *Fag.gra* | 0.62 | 23.7 | 33.2 | Y | Y | Y | Y | Y |
| *Fraxinus americana* | *Fra.ame* | 0.60 | 19.5 | 27.8 | Y | Y | Y | Y | Y |
| *Fraxinus nigra* | *Fra.nig* | 0.49 | 22.4 | 22.8 | Y | Y | Y | Y | N |
| *Fraxinus pennsylvanica* | *Fra.pen* | 0.56 | 17.9 | 31.2 | Y | Y | Y | Y | Y |
| *Fraxinus quadrangulata* | *Fra.qua* | 0.57 | 23.4 | 18.4 | N | N | Y | N | N |
| *Ilex opaca* | *Ile.opa* | 0.50 | 12.5 | 18.4 | Y | Y | Y | Y | Y |
| *Juglans cinerea* | *Jug.cin* | 0.38 | 18.8 | 26.6 | N | Y | Y | Y | N |
| *Juglans nigra* | *Jug.nig* | 0.54 | 22.5 | 40.2 | Y | Y | Y | Y | Y |
| *Juniperus virginiana* | *Jun.vir* | 0.49 | 4.1 | 22.1 | Y | Y | Y | Y | Y |
| *Larix laricina* | *Lar.lar* | 0.53 | 14.7 | 25.4 | Y | Y | Y | Y | N |
| *Liquidambar styraciflua* | *Liq.sty* | 0.48 | 19.0 | 32.7 | Y | Y | Y | Y | Y |
| *Liriodendron tulipifera* | *Lir.tul* | 0.36 | 27.8 | 52.8 | Y | Y | Y | Y | Y |
| *Maclura pomifera* | *Mac.pom* | 0.80 | 23.2 | 12.8 | Y | Y | Y | Y | N |
| *Magnolia acuminata* | *Mag.acu* | 0.48 | 22.4 | 28.1 | Y | Y | Y | Y | Y |
| *Magnolia fraseri* | *Mag.fra* | 0.44 | 22.5 | 23.4 | Y | Y | Y | Y | Y |
| *Magnolia grandiflora* | *Mag.gra* | 0.44 | 12.6 | 29.0 | N | Y | Y | N | N |
| *Magnolia virginiana* | *Mag.vir* | 0.45 | 15.7 | 19.5 | Y | Y | Y | Y | N |
| *Morus alba* | *Mor.alb* | 0.60 | 17.0 | 19.4 | N | Y | Y | N | N |
| *Nyssa sylvatica* | *Nys.syl* | 0.51 | 19.5 | 32.1 | Y | Y | Y | Y | Y |
| *Ostrya virginiana* | *Ost.vir* | 0.63 | 30.7 | 15.0 | Y | Y | Y | Y | Y |

**Table S10** continued.

|  |  |  | |  | |  | | Included in model? | | | | | | | | | |
| --- | --- | --- | --- | --- | --- | --- | --- | --- | --- | --- | --- | --- | --- | --- | --- | --- | --- |
| Species | Abbrev. | WD | | SLA | | HMax | | Sapling survival | | Canopy survival | | Growth | | Recruit- ment | | Repro. status | |
| *Oxydendrum arboreum* | *Oxy.arb* | | 0.54 | | 28.4 | | 19.7 | | Y | | Y | | Y | | Y | | Y |
| *Picea abies* | *Pic.abi* | | 0.47 | | 6.1 | | 50.1 | | Y | | Y | | Y | | Y | | N |
| *Picea glauca* | *Pic.gla* | | 0.41 | | 3.4 | | 29.1 | | Y | | Y | | Y | | Y | | Y |
| *Picea mariana* | *Pic.mar* | | 0.44 | | 4.2 | | 23.3 | | Y | | Y | | Y | | Y | | Y |
| *Picea rubens* | *Pic.rub* | | 0.39 | | 10.1 | | 32.8 | | Y | | Y | | Y | | Y | | Y |
| *Pinus banksiana* | *Pin.ban* | | 0.43 | | 5.8 | | 24.7 | | N | | Y | | Y | | N | | N |
| *Pinus echinata* | *Pin.ech* | | 0.51 | | 8.3 | | 38.7 | | Y | | Y | | Y | | Y | | Y |
| *Pinus elliottii* | *Pin.ell* | | 0.60 | | 5.3 | | 28.8 | | N | | Y | | Y | | N | | N |
| *Pinus glabra* | *Pin.gla* | | 0.43 | | 11.0 | | 28.2 | | N | | Y | | Y | | N | | N |
| *Pinus palustris* | *Pin.pal* | | 0.58 | | 3.6 | | 35.2 | | Y | | Y | | Y | | Y | | Y |
| *Pinus pungens* | *Pin.pun* | | 0.52 | | 7.6 | | 16.2 | | N | | Y | | Y | | Y | | N |
| *Pinus resinosa* | *Pin.res* | | 0.46 | | 4.9 | | 27.1 | | Y | | Y | | Y | | Y | | Y |
| *Pinus rigida* | *Pin.rig* | | 0.50 | | 5.9 | | 16.0 | | Y | | Y | | Y | | Y | | Y |
| *Pinus strobus* | *Pin.str* | | 0.36 | | 8.9 | | 48.5 | | Y | | Y | | Y | | Y | | Y |
| *Pinus sylvestris* | *Pin.syl* | | 0.43 | | 4.4 | | 39.7 | | Y | | Y | | Y | | Y | | N |
| *Pinus taeda* | *Pin.tae* | | 0.49 | | 10.4 | | 38.7 | | Y | | Y | | Y | | Y | | Y |
| *Pinus virginiana* | *Pin.vir* | | 0.49 | | 6.5 | | 23.0 | | Y | | Y | | Y | | Y | | Y |
| *Platanus occidentalis* | *Pla.occ* | | 0.42 | | 20.8 | | 34.4 | | Y | | Y | | Y | | Y | | N |
| *Populus balsamifera* | *Pop.bal* | | 0.33 | | 20.0 | | 29.7 | | Y | | Y | | Y | | Y | | N |
| *Populus deltoides* | *Pop.del* | | 0.43 | | 13.1 | | 40.5 | | N | | Y | | Y | | N | | N |
| *Populus grandidentata* | *Pop.gra* | | 0.39 | | 16.5 | | 21.2 | | Y | | Y | | Y | | Y | | N |
| *Populus tremuloides* | *Pop.tre* | | 0.36 | | 17.4 | | 17.5 | | Y | | Y | | Y | | Y | | Y |
| *Prunus avium* | *Pru.avi* | | 0.56 | | 18.4 | | 24.5 | | N | | Y | | Y | | Y | | N |
| *Prunus pensylvanica* | *Pru.pen* | | 0.36 | | 18.3 | | 19.6 | | Y | | Y | | Y | | Y | | Y |
| *Prunus serotina* | *Pru.ser* | | 0.50 | | 19.1 | | 34.4 | | Y | | Y | | Y | | Y | | Y |
| *Quercus alba* | *Que.alb* | | 0.65 | | 17.9 | | 37.4 | | Y | | Y | | Y | | Y | | Y |
| *Quercus bicolor* | *Que.bic* | | 0.67 | | 15.3 | | 25.9 | | N | | Y | | Y | | N | | N |
| *Quercus coccinea* | *Que.coc* | | 0.63 | | 14.1 | | 27.5 | | Y | | Y | | Y | | Y | | Y |
| *Quercus falcata* | *Que.fal* | | 0.61 | | 8.9 | | 23.1 | | Y | | Y | | Y | | Y | | Y |
| *Quercus laurifolia* | *Que.lau* | | 0.62 | | 12.5 | | 22.5 | | Y | | Y | | Y | | Y | | N |
| *Quercus macrocarpa* | *Que.mac* | | 0.63 | | 14.4 | | 30.9 | | N | | Y | | Y | | N | | N |
| *Quercus michauxii* | *Que.mic* | | 0.67 | | 12.7 | | 34.2 | | Y | | Y | | Y | | Y | | N |
| *Quercus nigra* | *Que.nig* | | 0.65 | | 10.9 | | 28.7 | | Y | | Y | | Y | | Y | | N |
| *Quercus palustris* | *Que.pal* | | 0.63 | | 13.5 | | 30.1 | | N | | Y | | Y | | Y | | N |
| *Quercus phellos* | *Que.phe* | | 0.64 | | 13.7 | | 30.0 | | Y | | Y | | Y | | Y | | Y |
| *Quercus rubra* | *Que.rub* | | 0.64 | | 15.0 | | 39.9 | | Y | | Y | | Y | | Y | | Y |
| *Quercus shumardii* | *Que.shu* | | 0.64 | | 11.8 | | 32.2 | | N | | Y | | Y | | Y | | N |
| *Quercus stellata* | *Que.ste* | | 0.71 | | 12.1 | | 21.5 | | Y | | Y | | Y | | Y | | Y |
| *Quercus velutina* | *Que.vel* | | 0.62 | | 14.6 | | 39.0 | | Y | | Y | | Y | | Y | | Y |
| *Robinia pseudoacacia* | *Rob.pse* | | 0.65 | | 26.6 | | 23.9 | | Y | | Y | | Y | | Y | | Y |
| *Salix nigra* | *Sal.nig* | | 0.39 | | 17.1 | | 30.8 | | Y | | Y | | Y | | Y | | N |
| *Sassafras albidum* | *Sas.alb* | | 0.46 | | 23.5 | | 22.7 | | Y | | Y | | Y | | Y | | Y |
| *Thuja occidentalis* | *Thu.occ* | | 0.31 | | 5.0 | | 17.8 | | Y | | Y | | Y | | Y | | Y |
| *Tilia americana* | *Til.ame* | | 0.36 | | 26.9 | | 34.6 | | Y | | Y | | Y | | Y | | Y |
| *Tsuga canadensis* | *Tsu.can* | | 0.41 | | 7.0 | | 32.0 | | Y | | Y | | Y | | Y | | Y |
| *Ulmus alata* | *Ulm.ala* | | 0.67 | | 22.7 | | 20.0 | | Y | | Y | | Y | | Y | | Y |
| *Ulmus americana* | *Ulm.ame* | | 0.51 | | 15.0 | | 36.6 | | Y | | Y | | Y | | Y | | Y |
| *Ulmus rubra* | *Ulm.rub* | | 0.52 | | 21.5 | | 27.9 | | Y | | Y | | Y | | Y | | Y |
